# Spatial modeling of telomere intra-nuclear distribution reveals non-random organization that varies during cell cycle and depends on LAP2 and BAF

**DOI:** 10.1101/2022.12.22.521599

**Authors:** Debora Keller, Sonia Stinus, David Umlauf, Edith Gourbeyre, Eric Biot, Nicolas Olivier, Pierre Mahou, Emmanuel Beaurepaire, Philippe Andrey, Laure Crabbe

## Abstract

Genome organization within the 3D nuclear volume influences major biological processes but is completely lost during mitosis, which represents a major challenge to maintain cellular identity and cell fate. To restore a functional G1 nucleus for the next cell cycle, it is imperative to reestablish genome organization during post-mitotic nuclear assembly. Importantly, the configuration of linear chromosomes has been shown to directly impact spatial genome architecture. Both centromeres and telomeres are known to associate with nuclear structures, such as the nuclear envelope, and support chromatin distribution. Here, using high-resolution 3D imaging combined with 3D spatial statistics and modeling, we showed that telomeres generally followed a regular distribution compared to what is expected under a random organization. While the preferential localization of telomeres at nuclear periphery was restricted to early G1, we found a strong clustering of centromeres in addition to their predominant peripheral localization at all cell cycle stages. We then conducted a targeted screen using MadID to identify the molecular pathways driving or maintaining telomere anchoring to the nuclear envelope. Among these factors, we could show that LAP2α transiently localizes to telomeres in anaphase, at a stage where LAP2α initiates the reformation of the nuclear envelope. Moreover, co-depletion of LAP proteins and their partner BAF impacted telomere redistribution in the next interphase. There results suggest that in addition to their crucial role in genome protection, telomeres also participate in reshaping functional G1 nuclei after mitosis.

## Introduction

The organization of the genetic material within the nucleus influences major biological processes, ranging from the regulation of gene expression to the timing of DNA replication and the maintenance of genome stability. As such, spatial genome architecture can impact cell fate and must be transmitted through cell lineages in order to maintain cellular identity. The nuclear envelope (NE), which defines the boundaries of the nuclear volume, confers the essential scaffold required to organize the nuclear content. The inner side of the NE is lined with a meshwork of intermediate filaments polymers forming the nuclear lamina, composed of A- and B-type lamins (Gruenbaum and Foisner, 2015). Together with additional NE-associated factors, the lamina provides a docking site for chromatin and creates a regulation hub for several essential cellular functions, including the establishment of the euchromatic active and heterochromatic inactive compartments within the nucleus, termed the A and B compartment, respectively (Falk et al., 2019; Solovei et al., 2016). This organization in specific compartments is visible as more condensed heterochromatic regions close to the NE, and the “lighter” more decondensed euchromatic compartments towards the center of the nucleus. Within these compartments, many local and long-range contacts further organize chromatin in loops and Topologically Associated Domains (TADs).

How 3D organization is established and what drives chromatin segregation by scaffolding is still unclear. Recent evidence suggests that centromeres, whose primary function is to promote proper chromosome segregation during cell division, could directly impact spatial genome architecture (Muller et al., 2019). Many studies in the budding yeast *Saccharomyces cerevisiae* and fission yeast *S. pombe* have clearly established a strong clustering of centromeres all along the cell cycle (Funabiki et al., 1993; Guacci et al., 1997; Jin et al., 2000), feature that is conserved in higher eukaryotes (Solovei et al., 2004; Vourc’h et al., 1993; Weierich et al., 2003). Centromere clustering impacts chromatin organization by creating a subcompartment within the nucleus, as well as a barrier to intrachromosomal arm interactions. Importantly, a frequent localization of centromere clusters toward the nuclear periphery was also reported (Muller et al., 2019). In yeast, centromeres cluster in one focus near the spindle pole body, opposite the nucleolus, with a strong impact on genome spatial regulation. Studies in mouse and human also pointed to a localization of centromeres at the nuclear periphery. In human lymphocytes, 62% of pericentromeric heterochromatin was found in the B compartments associated with the lamina and the nucleolus (Weierich et al., 2003).

Similar to centromeres, the nuclear position of telomeres, ribonucleoprotein complexes located at the ends of linear chromosomes, has also been intensely studied in yeast, and has influenced the search for potential connections of mammalian telomeres with nuclear structures. In *S. cerevisiae*, telomeres are organized as clusters tethered to the inner nuclear membrane (INM) via the SUN-domain protein Mps3 (Conrad et al., 2007), which promotes telomere silencing and the inhibition of unwanted recombination events between telomeric repeats (Hediger et al., 2002; Schober et al., 2009). Telomere dynamics also play an essential function during meiosis in budding yeast, fission yeast, and in mice, as their clustering to the nuclear envelope is crucial for proper meiotic pairing and recombination of homologous chromosomes (Chikashige et al., 2009, 2007, 2006, 1994). By contrast, nuclear distribution of human telomeres is still very elusive. The dynamic behavior of human telomeres was previously studied over a period of a few hours, which uncovered their constrained diffusive movement, and the formation of dynamic clusters (Molenaar et al., 2003). 3D-fluorescent in situ hybridization (FISH) confocal microscopy experiments performed on human lymphocytes showed that telomeres are on average nearer to the center of the cell than centromeres and are not enriched at the nuclear periphery in interphase cells (Amrichová et al., 2003a; Weierich et al., 2003). A similar method followed by quantitative analysis to determine nuclear telomeric organization on fixed mouse and human lymphocytes established that telomeres assemble into a telomeric disk specifically in the G2 phase, suggesting a cell cycle regulation aspect of telomere position (Chuang et al., 2004). Cell cycle regulation of telomere positioning was later evidenced by spinning-disk confocal timelapse microscopy over an entire cell cycle. It was uncovered that ~45% of human telomeres are physically attached to the NE during postmitotic nuclear assembly, both in primary fibroblasts and cancer cells (Crabbe et al., 2012). However, distance analyses were performed in 2D, using the middle plane of the cell, which corresponds only to a representative fraction of all telomeres within the 3D nuclear volume.

Mitosis represents one of the greatest challenges to maintain cellular identity. While chromatin is condensed up to 50-fold in metaphase chromosomes, TAD and chromosome compartments are lost at this stage (Naumova et al., 2013). In addition, NE breakdown that characterizes open mitosis totally resets nuclear structure. During post-mitotic nuclear assembly, nuclear size needs to be readjusted, nuclear pores that allow trafficking between the nucleus and the cytoplasm reinserted within the NE, and chromosome territories established (Güttinger et al., 2009). These events must be finely coordinated to ensure that segregated DNA is finally enclosed in a single cell nucleus in each daughter cell. During late anaphase/telophase, INM proteins bind chromatin to initiate attachment of membrane sheets (Haraguchi et al., 2008). Early live microscopy studies using fluorescently-tagged proteins expressed in HeLa cells found that chromosome ends associate transiently to the INM proteins Lamina-Associated Polypeptide (LAP) 2α and Barrier-to-Autointegration Factor (BAF) (Dechat et al., 2004), suggesting a role of telomeres in chromatin reorganization at this stage. These results are in accordance with the observation that telomeres decorate the NE in early G1, where they are found transiently interacting with lamins, LAP2α, and Emerin specifically during postmitotic nuclear reformation (Crabbe et al., 2012).

While the above-mentioned studies revealed a link between nuclear re-organization and telomeres, to our knowledge, there is no systematic analysis of telomere spatial positioning within the nucleus and across the cell cycle that could shed light on the role of telomeres in nuclear organization. To address this issue, we used a pipeline combining high-resolution 3D imaging of telomeres and NE proteins at specific cell cycle stages, complemented with novel 3D spatial statistics and modelling (Arpòn et al., 2021). We found that telomeres are organized in a non-random fashion in the nucleus and are undergoing a dynamic repositioning from the periphery to the interior as cells progress from early G1, through G1/S and G2. By contrast and consistent with previous work in other cell lines, centromeres remained predominantly associated with the periphery in a polarized way. We then conducted a targeted screen using MadID, a proximity labeling approach to map protein-DNA interaction (Sobecki et al., 2018; Umlauf et al., 2020), that revealed a set of factors involved in telomere tethering to the NE. Among these factors, we further showed that under endogenous conditions LAP2 proteins associated with telomeres in anaphase, at the onset of NE reformation. Co-depletion of BAF and LAP proteins affected the size of reforming nuclei after mitosis, and the nuclear distribution of telomeres in the subsequent interphase.

## Results

### 3D Structured Illumination Microscopy (3D-SIM) to assess telomere and centromere spatial organization

To study the organization of telomeres and centromeres within the nucleus at given cell cycle stages and with high 3D resolution, we turned to 3D structured illumination microscopy (3D-SIM), as it allows rapid multi-color imaging over the depth of a cell at 8-fold increased volumetric resolution over the diffraction limit (Schermelleh et al., 2019). To obtain a high signal-to-noise ratio for telomere segmentation, we used HeLa cells expressing EGFP-TRF1, one of the core proteins from the Shelterin complex that sits on telomeric repeats. Telomere capping and cell cycle progression were previously shown to be unaffected in EGFP-TRF1 overexpressing HeLa cells (Crabbe et al., 2012). Centromeres were stained using CREST and the nuclear envelope was visualized by co-staining of LaminA/C, a component of the nuclear lamina, and SUN1, a transmembrane protein localized at the INM. Cells were fixed for immunofluorescence following synchronization at the G1/S boundary, in G2 phase, or in late mitosis/early G1 phase (Fig. S1A-B). Early G1 cells that are still in the process of postmitotic nuclear assembly display the typical SUN1 aggregates around the nucleus, corresponding to the portion of the NE protein not yet localized at the envelope (Fig. S1B) (Crabbe et al., 2012). Early G1 cells readily reach sizes of about 15-20 μm, thus often leading to decreased 3D-SIM image contrast. To circumvent this effect, we devised a 3D-SIM mounting medium based on the previously reported clearing agent sorbitol (Szczurek et al., 2018a), and assessed 3D-SIM modulation contrast by SIMCheck (Ball et al., 2015a) and reconstruction quality (Fig. 1A-S1C) prior to analysis. We quantitatively assessed in 3D various morphometric descriptors based on a specifically devised image processing and segmentation pipeline, including an echo-suppression algorithm (Fig. 1A and Methods section). Segmentation was performed on images of individual nuclei from two datasets stained for nuclear lamina and either telomeres or centromeres.

**Fig. 1.**
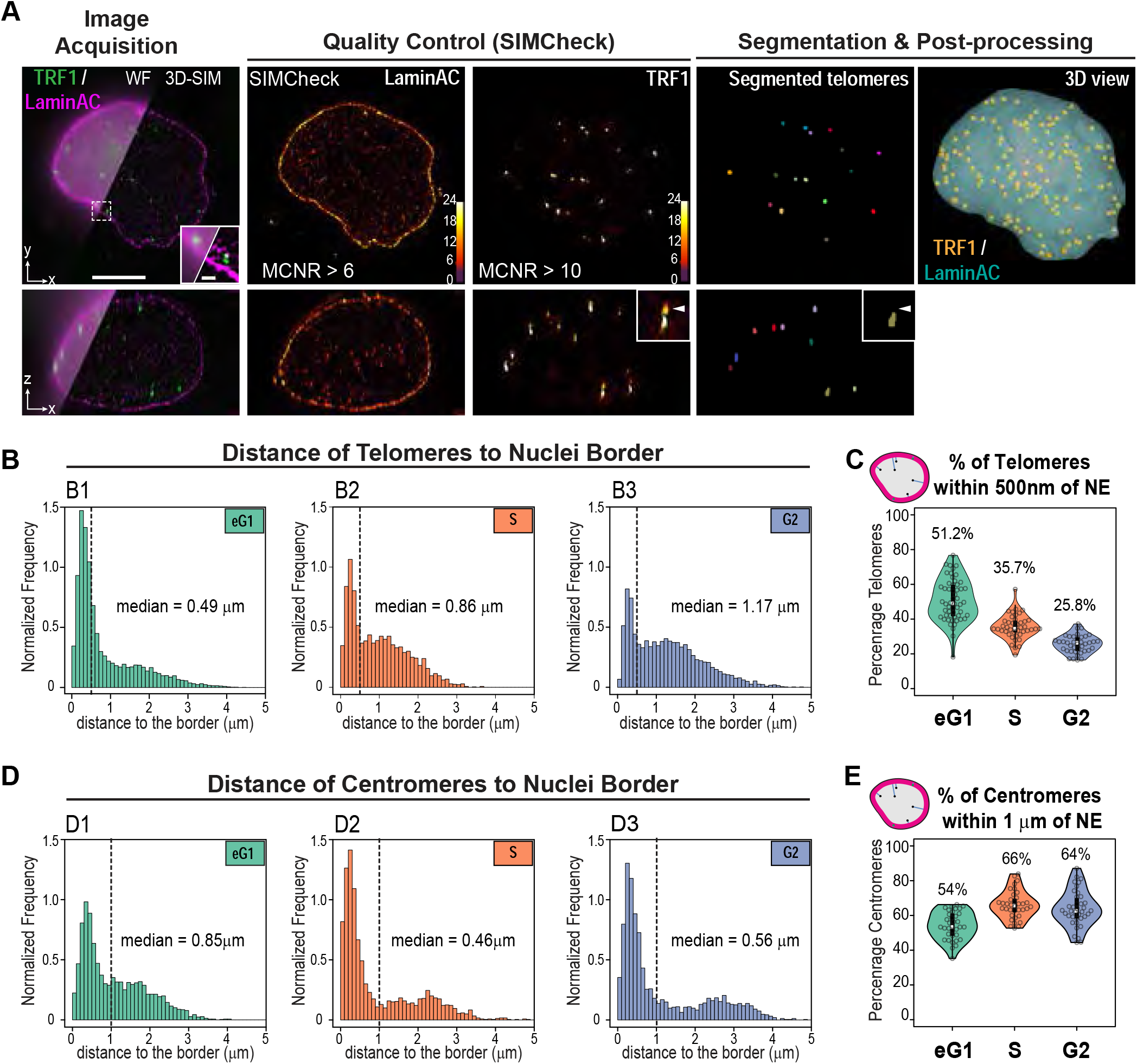
Quantitative analysis of 3D-SIM images of HeLa cell nuclei shows dynamic positioning of telomeres during cell cycle. A. Overview of the analysis pipeline from image acquisition to segmentation & post-processing. Representative images of nuclei stained for TRF1 (*green*) and LaminAC (*magenta*) highlighting the increased resolution in 3D-SIM compared to pseudo widefield (WF, top left) in the lateral *xy* and axial *xz* directions. The quality of 3D-SIM images was assessed using SIMCheck (Ball et al., 2015b), cells with adequate modulation contrast (MCNR) values for LaminAC and TRF1 are then segmented and processed (see also **Figs. S1C**). Telomeres segmented with a specifically developed pipeline described in the Methods section are shown in individual XY and XZ sections. *Inset:* artefactual echoes in the SIM reconstruction are removed by the segmentation pipeline. 3D view shows segmented telomeres (*Yellow dots*) within a segmented nucleus (*Blue surface*). Scale bars: 5 μm (main) and 1 μm (inset). B. Distribution of distances between telomeres and nuclear border during cell cycle and their median value showing increased distance to the NE (*B1:* early G1 phase, N = 54 nuclei; *B2:* S phase, N = 43 nuclei; *B3:* G2 phase, N = 39 nuclei). Distance was measured from the center of each telomere to the closest point at the nuclear surface. Frequencies were normalized to obtain a unit area histogram. C. Distribution of percentage of telomeres located at a distance below 500 nm from nuclear border in individual nuclei during cell cycle. D. Same as B for centromeres (N = 35, 36 and 38 for early G1, S and G2 phases, respectively). E. Distribution of percentage of centromeres located at a distance below 1 μm from nuclear border in individual nuclei during cell cycle.

For both datasets, we confirmed that the nuclear morphology varied across the cell cycle (Fig. S1D-E). The volume of late mitosis and early G1 nuclei exhibited a bimodal distribution, corresponding to the transition from telophase (< ~800 μm^3^) to a more decondensed state where nuclei are bigger (> ~800 μm^3^). Nuclear volume almost doubled when transitioning from early G1 to G2 (from 800 μm^3^ G1/S nuclei to 1300 μm^3^ G2 nuclei). Centromere size and shape also varied during the cell cycle, with volumes ranging from 0.21+/-0.01 μm^3^ in early G1 to 0.27+/-0.01 μm^3^ in G2 (Fig. S1E). By contrast, the number of segmented centromeres decreased from ~90 in early G1 to ~50 in S and ~60 in G2. The concomitant decrease of centromere number and increase of their volume points to centromere clustering that cannot be resolved in 3D-SIM, also observed in previous studies (Solovei et al., 2004; Weierich et al., 2003), or reflect the structural evolution of the centromeric complex as described for CENP-A, transitioning from a globular rosette in eG1 to a less structured and wider disc in early mitosis (Andronov et al., 2019). Telomere number also varied throughout the cell cycle (Fig. S1D). We segmented 142±2 telomeres per nucleus in early G1, which corresponds to the ~70 chromosomes described in HeLa cells. The number of telomeres decreased to 65±2 and 91±3 in G1/S and G2 phases, respectively. This suggests that telomeres also undergo clustering, as previously observed (Adam et al., 2019; Chuang et al., 2004; Crabbe et al., 2012; Nagele et al., 2001). Concomitant to the increase in nuclear volume, this trend resulted in an overall decrease of telomere density between early G1 and G2. Overall, our results confirmed that 3D-SIM combined with specific image analysis pipelines was a valid approach to quantify in 3D and with a high-resolution the dynamics of telomeres and centromeres during the cell cycle.

### Spatial statistical modeling shows dynamic positioning of telomeres relative to the NE over the cell cycle while centromeres remain stably associated

To study telomere distribution, we first analyzed the distance of each telomere to the edge of the nucleus in early G1, G1/S and G2 cells (Fig. 1B). We found a bimodal distribution in all three cell cycle phases, with a pool of telomeres localized within 500 nm of the NE (dotted line) – the estimated thickness of the LADs contacting the nuclear lamina (Kind et al., 2015) – and a population of more internally located telomeres with a larger spread of distances to NE. In accordance with a previous study using 2D image analysis (Crabbe et al., 2012), more than 50% of telomeres were found within 500 nm of the NE in early G1, with a median distance to the NE of 0.49 μm (Fig. 1B-C). The proportion of these peripheral telomeres gradually decreased during the cell cycle, to reach ~36% and ~26% of telomeres within 500 nm of the NE in G1/S and G2, respectively. These results suggest the existence of two sub-populations of telomeres that undergo a dynamic switch between the periphery and the nuclear interior as cells progress through the cell cycle, with only ~20 telomeres that remained near the NE (Fig. S1D). Centromeres also exhibited a bimodal distribution of distance to the nuclear border, with a cutoff between peripheral and internal centromeric sub-populations at ~1 μm of the border (Fig 1D, dotted line), likely due to the 10-fold larger size of centromeres compared to telomeres and accompanying steric constraints. However, unlike telomeres, the proportion of peripheral centromeres increased from 54% in early G1 to 66% in G1/S and remained stable at 64% in G2 (Fig 1E). Absolute sub-population sizes showed this trend was due to a decrease in the number of internal centromeres at G1/S, while the number of peripheral centromeres remained stable throughout the cell cycle (Fig. S1E), indicating a more stable proximity to the NE.

The radial positioning of chromatin is functionally relevant, as the nuclear periphery is known to be a domain boundary that regulates chromatin function and organization. However, though they provide valuable information, distance measurements are not sufficient to assess spatial proximity with nuclear border. For example, about 90% of the volume in a sphere is closer to the border than to the center (Arpòn et al., 2021a). Therefore, we adopted a statistical spatial modeling approach recently developed to assess 3D spatial interactions in object patterns (Arpòn et al., 2021a). Here, spatial interactions mean interdependent positioning of objects (for example attraction or repulsion), which does not necessarily imply direct physical interactions. In this method, the observed 3D spatial distribution of objects within a single nucleus is quantitatively described using distance functions (detailed below) and compared to a theoretical distribution model of those same objects within this same nucleus (Fig. 2A). The difference between the observed pattern and model predictions is quantified by a Spatial Distribution Index (SDI) that varies between 0 (observed distances far below predictions) and 1 (observed distances far above predictions). At the population level, the model is rejected if the SDI distribution differs from the uniform distribution between 0 and 1 (Fig. 2A). We first examined the distance of objects to the nuclear periphery using the B function (Fig. S2A) and compared observations to the completely random model, in which objects are uniformly and independently distributed within the nuclear space. We found that in early G1 nuclei, the positioning of telomeres relative to the periphery strongly deviated from the random model, with smaller distances to the nuclear boundary than expected under randomness (Fig. S2B). This positive spatial interaction with NE in early G1 was confirmed at the population level with a skewed distribution of B-SDI values towards 0 (Fig. 2B). Preferential association of telomeres with the periphery was lost once cells were at the G1/S boundary (Fig. 2B). Analysis of G2 nuclei showed an inverted pattern of interactions with NE compared with early G1 nuclei, with telomeres exhibiting larger distances from the NE than under the random model. The C function, which probes the distance of objects to the centroid of nuclei (Fig. S2A), corroborated these results and the switch from globally positive to globally negative spatial interactions between telomeres and NE between early G1 and G2 (Fig. S2C). Taken together, our analysis revealed a switch in the radial positioning of telomeres and in telomere-NE spatial interactions during the cell cycle, with first a decrease in the number of peripheral telomeres followed by an increase in the number of internal telomeres as cells progress from early G1 to G2. Centromeres behaved very differently and exhibited a consistently strong and positive nonrandom spatial interaction with the nuclear periphery that was maintained throughout the cell cycle, and even reinforced from G1/S with the decrease in the number of internally located centromeres (Fig. 2C & S2D-E).

**Fig. 2.**
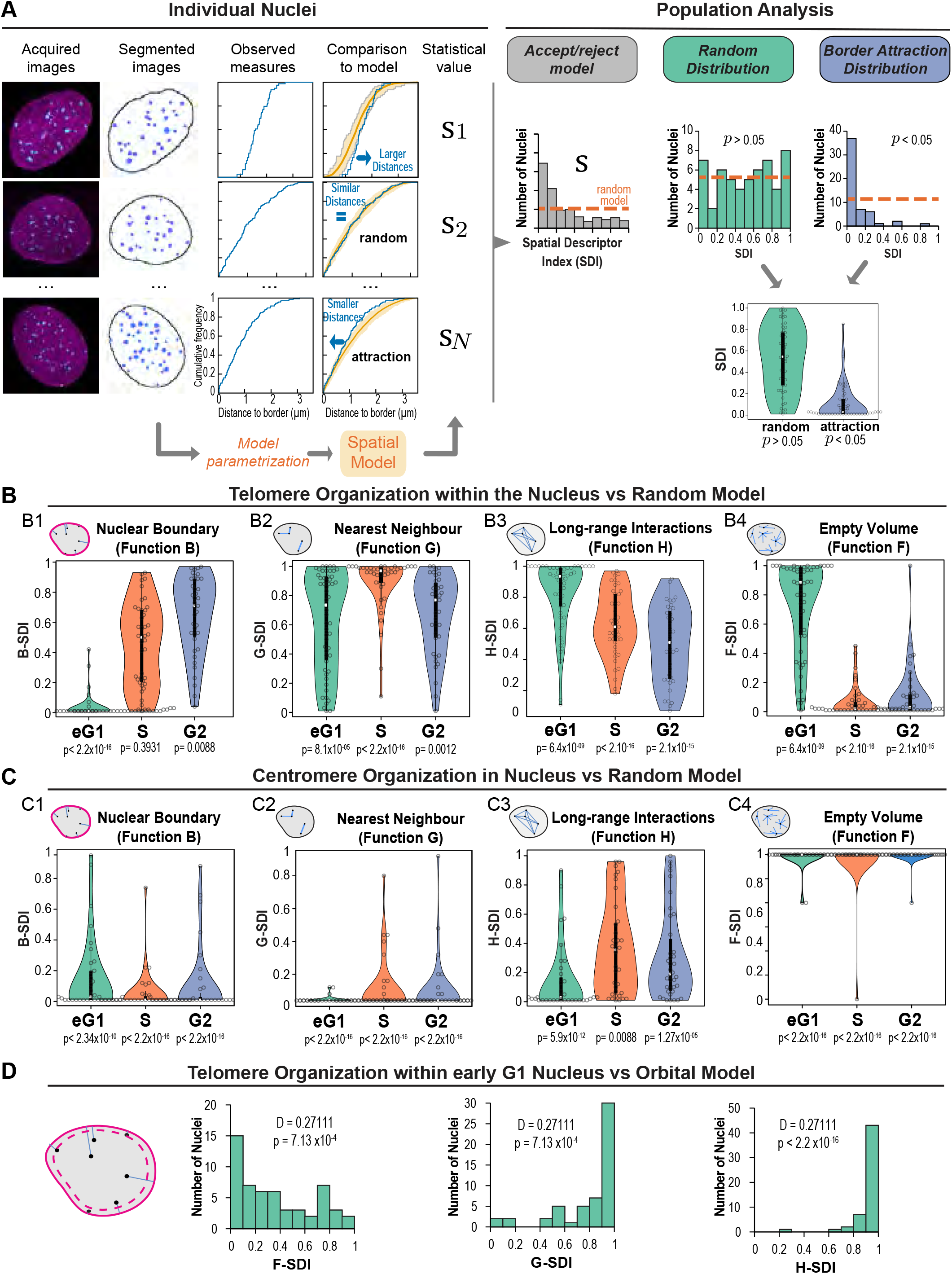
Statistical spatial analysis shows a dynamic switch in the spatial interactions between telomeres and nuclear envelope during the cell cycle. A. Statistical spatial analysis pipeline: illustration with the testing of spatial interaction with the nuclear border. Based on the cumulative distribution function (CDF) of the distance between telomeres and nuclear border, observed individual patterns were compared to predicted patterns under a random model of organization. For each pattern, the probability of observing smaller distances under the model than actually observed was computed (S=Spatial Descriptor Index, SDI). Upon a positive spatial interaction (attraction) between telomeres and nuclear border, small values of the SDI are expected because smaller distances should be observed as compared to model predictions. In the absence of any spatial interaction, a uniform distribution is expected (*Orange dotted line*). The random model is tested by testing the uniformity of the population distribution of the SDI. B. Analysis of spatial interactions between telomeres and nuclear border (*B1*) and between telomeres (*B2-B4*) using comparisons to the random model of telomere organization. *B1*: distribution of SDI computed using the CDF of the distance between each telomere and nuclear border (p: p-value of Kolmogorov-Smironov test of uniformity). *B2-B4:* same as (B1) for SDIs computed based on the cumulative distribution function of the distance between each telomere and its closest neighbor (*B2*), of the distance between each telomere and any other telomere (*B3*), and of the distance between arbitrary nuclear positions and their closest telomeres (*B4*). Early G1 phase, N = 54 nuclei; S phase, N = 43 nuclei; G2 phase, N = 39 nuclei. C. Same as B for centromeres. Early G1 phase, N = 35 nuclei; S phase, N = 36 nuclei; G2 phase, N = 38 nuclei. D. Analysis of spatial interactions between telomeres in early G1 using comparisons to the orbital model of telomere organization (N=54 nuclei). The scheme illustrates the orbital model, which is similar to the completely random model of telomere organization except that the observed distance between each telomere and nuclear border is preserved.

### Polarity in telomere and centromere distribution

We took a closer look at the 3D organization of both telomeres and centromeres and observed a difference in polarity along the minor axis of the nucleus. Early G1 telomeres were predominantly found on one side of the nucleus while centromeres were located on the opposite side (Fig. S3A-B), which is likely a consequence of the pulling forces exerted on centromeres during anaphase that lead to a Rabl-like configuration. We therefore quantitatively assessed the polarity of telomeres and of centromeres along each of the three main axes of the nucleus and defined a polarity index based on the proportion of positions that projected on each half-axis (Fig. S3C). These polarity indexes confirmed the early G1 asymmetric distribution of both telomeres and centromeres along the minor axis of the nucleus and the absence of polarity along the two other axes (Fig. S3D and E). However, while telomeres lost their polar organization in later stages, centromeres exhibited a sustained high polarity index. These data suggest that a Rabl-like configuration of centromeres is, to some extent, preserved during the whole cell cycle. This observation could not be made based on the 2D analysis of the middle plane of the cell (Fig. S3A-B, dotted line) that was initially performed (Crabbe et al., 2012). Overall, our results stress the importance of coupling 3D high-resolution imaging and quantitative image analysis to unbiased statistical schemes that assess spatial interactions at the population level despite heterogeneity in nuclear morphology and in the number and sizes of nuclear objects.

### Telomeres exhibit a multi-scale organization in the nuclear volume in early G1

We next asked whether additional spatial interactions were subtending telomere organization. We relied on the same spatial modeling approach as above, comparing observed patterns to predictions of the random distribution model using distance functions that probe the relative positioning between objects. We first analyzed the distance between each object and its nearest neighbor, using the function G (Fig. S2A). In early G1, telomeres showed a mixed behavior with an attractive trend in the short distance range (shorter distances to nearest neighbor than under the random model at distances < ~1 μm) and a repulsive trend in the long-distance range (larger distances than under the model at distances > 1 μm) (Fig S4A-B). This mixed response was reflected at the population level by the bimodal distribution of the corresponding G-SDI (Fig. 2B). The long-range regularity was confirmed by analyzing the distances between all pairs of telomeres (function H; Fig. S2A), which showed on average larger distances than expected under a random organization, both on individual patterns (H-H0>0 at distances > 2 μm – Fig. S4C) and at the population scale (Fig. 2B). Function F, which probes the empty spaces between objects (Fig. S2A), showed that at both the individual and the population levels the voids between telomeres were larger than expected under randomness (Figs. S4D, 2B), thus confirming the trend for clustering in the short-range scale. Overall, the three different spatial descriptors suggest a multi-scale organization of telomeres in early G1, with telomeres being excluded from large nuclear domains yet showing a regular distribution in the domains they occupy.

### Telomere organization is subtended by mutually repulsive spatial interactions

We wondered whether the preferentially peripheral positioning of telomeres in early G1 was sufficient to explain the short-range attraction or, conversely, the long-range repulsion. We evaluated these hypotheses by comparing observed patterns to predicted distributions under the orbital model (Arpòn et al., 2021). This model enforces identical relative positioning to the nuclear border in both observed and predicted patterns. The SDI-distributions obtained with the three distance functions all pointed to a more regular distribution of telomeres than predicted under a purely orbital distribution and showed no tendency for clustering (Fig. 2D). This shows that in the comparison to the completely random model above, the demonstrated attractive effect was probably due to the clustering of telomeres at the nuclear periphery while the repulsive effect could not. Hence, the orbital model confirms that telomere organization is not only radial but is also subtended by mutually repulsive spatial interactions.

As cells progressed through the cell cycle, the mixed behavior observed in early G1 with both attraction and repulsion was lost. In S/G1, observed distances to nearest neighbor were larger than expected under the random model (Figs. S4B, 2B2). The loss of the attractive trend was confirmed by analysis using functions F and H, which showed more regular and smaller voids between telomeres and larger inter-distances between telomeres, respectively, than expected under a random model, thus pointing to a regular distribution (Figs. S4C-D, 2B3-4). Functions F and G showed regularity was maintained in G2 at least in the short distance range, since the observed H-SDI distribution did not differ from complete randomness (Fig 2B2, 2B4). Thus, at the G1/S transition and G2, telomeres exhibited a purely repulsive and regular organization, devoid of clustering. Overall, telomeres followed throughout the cell cycle a more regular distribution than expected under a random organization, a possible consequence of the partition of the nucleus into chromosome territories. In early G1, a clustering effect superimposed onto this repulsive pattern, likely due to the peripheral polar organization of telomeres along the nucleus minor axis.

From early G1 to G2, distances between each centromere and its nearest neighbor were consistently smaller than expected under a random model (Fig. 2C2), thus showing a preserved trend for a clustered organization. This attractive pattern was confirmed by the larger spaces devoid of centromeres evidenced by function F and by the globally smaller inter-distances between centromeres shown by function H (Figs. 2C3, 2C4). Taken together, our spatial analyses confirm a strong clustering of centromeres in addition to their predominantly peripheral localization within the nucleus at all cell cycle stages and show a more stable organization throughout the cell cycle than observed for telomeres.

### Targeted screen for factors involved in telomere-NE anchoring

Next, we aimed to identify the actors involved in 3D-telomere organization and in telomere tethering to the NE. We took advantage of MadID, a tool we recently developed to probe for protein-DNA interactions *in vivo* using proximity labeling (Sobecki et al., 2018; Umlauf et al., 2020). MadID relies on the expression of the bacterial methyltransferase M.EcoGII, which adds methyl groups to N6-adenosine (m6A) in any DNA sequence context. When fused to LaminB1, M.EcoGII specifically methylates chromatin that comes in contact to the nuclear lamina. This technique was previously used to map Lamina Associated Domains (LADs) by whole genome sequencing with high specificity, and deep and unbiased genome coverage (Sobecki et al., 2018). MadID could also probe telomeres-NE contact sites in human cells, which we decided to use as readout for a siRNA-mediated targeted screen. We selected 34 targets, either because of their suspected role in chromatin organization, or after a mass spectrometry screen we performed to uncover hits involved in telomere-NE anchoring (unpublished work). These factors could be classified in four different groups: i) members of the shelterin complex and related proteins; ii) members of the Linker of Nucleoskeleton and Cytoskeleton (LINC) complex; iii) members of the nuclear pore complex (NPC); iv) proteins from the NE and related (Fig. 3A). HeLa cells were transduced with inducible retroviral vectors expressing either M.EcoGII-LaminB1 (M-LB1), or unfused M.EcoGII that was used as a reference to correct for local differences in chromatin accessibility. Clonal cell populations expressing equal levels of M.EcoGII or M-LB1 were isolated. Following 24 hours of Shield-1-dependent induction, M-LB1 was found properly localized at the nuclear rim (Fig. S5A), and catalyzed genomic DNA methylation detected with an m6A-specific antibody *in situ* by DNA-IF (Fig. S5B) and immuno-dot-blot (Fig. S5C) (Sobecki et al., 2018; Umlauf et al., 2020). To test whether MadID could be performed in these clonal cells with high specificity and reproducibility, we performed m6A-specific immunoprecipitations (m6A-IP) followed by quantitative PCR (qPCR) using primers specific to the well-established LAD-CFHR3 and inter-LAD-SMIM2 regions (iLAD) (Kind et al., 2013), as well as to telomeric repeats (Kychygina et al., 2021a). We found a 30-fold enrichment of LAD-CFHR3 and a 10-fold enrichment of telomeric repeats specifically in M-LB1 cells, further confirming the proximity of a subset of telomeres to the nuclear envelope (Fig. 3B). By contrast, HeLa cells expressing M.EcoGII fused to the shelterin protein TRF1 (M-TRF1) to directly address the methyltransferase to telomeres (Sobecki et al., 2018) triggered methylation of telomeric repeats (Fig. 3B) but not of the LAD-CFHR3 region (Fig. 3B).

**Fig. 3.**
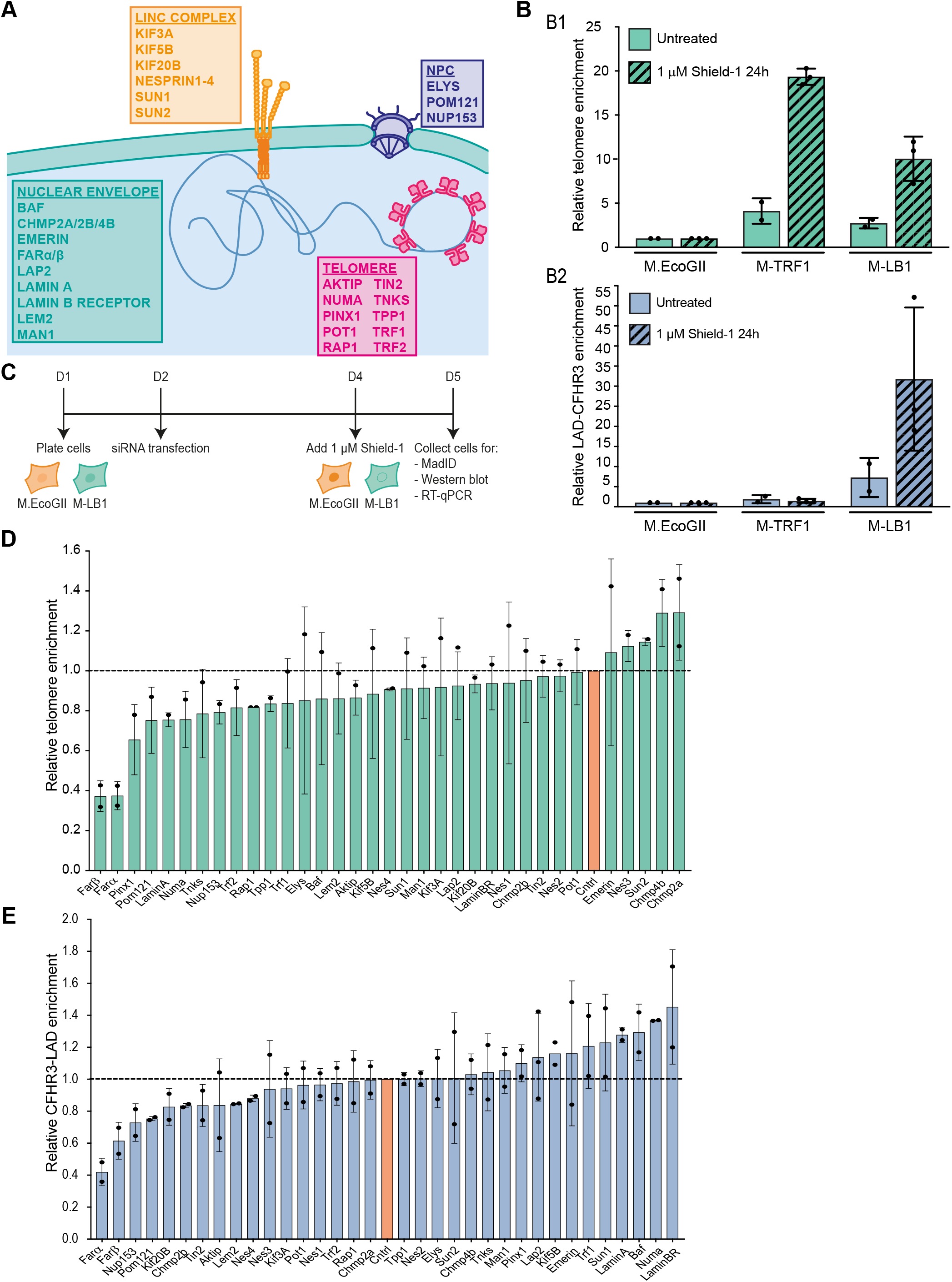
MadID-based targeted screen reveals factors involved in telomere interaction with the nuclear envelope. A. Scheme of proteins selected for the targeted MadID screen. Selected proteins belong to four categories: members of the linker of nucleoskeleton and cytoskeleton complex (LINC, orange), members of the nuclear pore complex (blue), members of the shelterin complex (magenta) and nuclear envelope-related proteins (green). B. Experimental setup of MadID-based targeted screen. C. Relative telomere (top) and LAD-CFHR3 (bottom) enrichment in cells expressing M.EcoGII, M-TRF1 or M-LB1, in absence (n=2) or presence (n=3) of Shield1 (1 μM, 24h). Mean ± SD is shown. D. Relative telomere enrichment upon depletion of the indicated proteins. Telomere enrichment is shown relative to iLAD-SMIM2 and to control condition (set at 1). n=2 except for LAP2 (n=3). Mean ± SD is shown. E. Relative LAD-CFHR3 enrichment upon depletion of the indicated proteins. LAD-CFHR3 enrichment is shown relative to iLAD-SMIM2 and to control condition (set at 1). n=2 except for LAP2 (n=3). Mean ± SD is shown.

After validation of the clonal cell populations, we implemented the screen following the experimental set-up as shown in Fig. 3C. Briefly, clonal cells were transfected with siRNA for 72 hours, and M.EcoGII or M-LB1 expression was induced by addition of Shield-1 during the last 24 hours before collection. Efficiency of target depletion could be verified for 32/34 the candidates by western blotting or RT-qPCR, with 2 targets for which no specific antibodies or good PCR primers were found (Fig. S6.1-6.2). After genomic DNA extraction, m6A-IP followed by qPCR was performed to reveal telomeric and LAD enrichments in the IP fraction. Importantly, cycle threshold values for all three regions as well as the relative fold enrichments over SMIM2 obtained for siControl replicates were highly reproducible across multiple independent siRNA experiments (Fig. S5D-E). We then plotted the relative telomere and LAD-CFHR3 enrichments for all tested siRNA relative to siControls (Fig. 3D-E). The obtained graph indicates whether the depletion of each target decreased (values below 1) or increased (values above 1) contact frequencies between LAD-CFHR3 or telomeric repeats and LaminB1. To further assess the specificity of our screen, siRNA against farnesyltransferase (FNT) alpha and beta were used to perturb LaminB1 localization at the NE. A- and B-type lamins are major farnesylated proteins, and while the mature form of LaminA loses its farnesylated site by cleavage during its final processing, LaminB1 and LaminB2 keep their farnesyl moiety. We therefore postulated that siRNA against FARα or FARβ could prevent proper addressing of M-LB1 to the NE, similarly to the addition of farnesyl transferase inhibitors that mislocalize B-type lamins to the nucleoplasm (Adam et al., 2013). Indeed, we found a strong defect in M-LB1 addressing to the NE in cells depleted for either FNT (Fig. S5F-G), and a 40 to 60% reduction in LAD-CFHR3 and telomeres enrichment in the m6A-IP fraction (Fig. 3D-E).

Overall, we found that most selected targets had a positive effect on telomere tethering, as their depletion reduced telomere enrichment in the m6A-IP fraction compared to siControls (Fig. 3D). Hence, depletion of members of the shelterin complex decreased contacts with the NE, suggesting that telomeric proteins engage, to some extent, in protein-protein interactions leading to telomere tethering. Except for TIN2, depletion of members of shelterin did not affect LAD-CFHR3 contacts (Fig. 3E). Depletion of members of the LINC complex, such as SUN1, kinesins and nesprins, also altered telomeres-NE contact sites. This result is coherent with the role of the LINC complex in telomere tethering in several organisms, as well as its role in chromatin movement during meiosis (Link et al., 2015). A similar trend was found for the members of the NPC complex POM121, ELYS, and NUP153, which were previously shown to be involved in the regulation of genome architecture or NE reformation (Anderson et al., 2009; Haraguchi et al., 2000; Lemaitre et al., 2012; Ptak and Wozniak, 2016; Shevelyov, 2020; Talamas and Hetzer, 2011). In contrast, members of the ESCRT-III complex CHMP4B and CHMP2A seemed to prevent telomere anchoring, as their depletion increased telomere-LaminB1 interaction. Described for its role in cytoplasmic membrane fusion, the ESCRT-III complex is also known to remodel the NE, particularly during mitotic exit (Olmos et al., 2015; Vietri et al., 2015). CHMP2A is recruited to the reforming NE in late anaphase through CHMP4B, providing an essential function for NE reformation. Importantly, CHMP2B, which did not trigger a change in telomere methylation status, has minimal effect on NE reformation by itself, and requires co-depletion with CHMP2A to trigger a greater phenotype (Olmos et al., 2015). Among other proteins associated with the NE, LaminA and BAF or LAP proteins had an interesting effect on chromatin attachment to the NE. Indeed, their depletion increased the interaction between LAD-CFHR3 and LaminB1, as previously shown (Kind and Steensel, 2014), but decreased telomere attachment (Fig. 3D-E).

### Lap2α associates with telomeres in anaphase

While our targeted screen provided a first hint on the molecular pathways driving or maintaining telomere-NE attachment in human cells, we sought to address the establishment of such interaction, which takes place at the end of mitosis. In late anaphase, membranes need to reform around chromatin, and evidence suggests that endoplasmic reticulum membrane tubules are targeted to chromatin through chromatin-binding NE proteins (Anderson and Hetzer, 2008; Kutay and Hetzer, 2008). The major NE protein association sites on chromatin, called peripheral or core regions, attract different sets of proteins by poorly understood mechanisms. A pioneer study using overexpression of fluorescently tagged proteins showed that LAP2α transiently localizes in anaphase to telomeres at inner core regions to initiate NE reformation (Dechat et al., 2004). A subfraction of BAF is also relocalized to core structures together with LAP2α. These results suggest that telomere-NE anchoring observed during postmitotic nuclear assembly (Crabbe et al., 2012) could initiate in anaphase, and that telomeres could serve as a nucleation point for membrane reformation. To test this hypothesis, we labeled endogenous LAP2α by immunofluorescence during all stages of mitosis and acquired images using confocal microscopy followed by deconvolution (Fig. 4A). As expected, endogenous LAP2α was found mainly diffuse in the nucleoplasm during interphase, and transiently recruited at core regions of chromatin early in anaphase, with accumulation of the protein at discrete sites that often overlapped with telomeric signal (Figs. 4A & S7A). Instead, LAP2β isoform was rather observed at peripheral regions before the signal extends all around chromatin in later mitotic stages (Fig. 4A, S7A). Following image segmentation, we quantified the distance between the centroid of telomeres and the edge of LAP2α/β labeled regions using BIP software (see Material and Methods). About 75% of telomere foci in anaphase and 82% in telophase were found within 200 nm of LAP2 structures (Figs. 4B, S7A). Altogether, these results suggest that NE proteins involved in membrane reformation are recruited at or close to telomeres during postmitotic nuclear assembly.

**Fig. 4.**
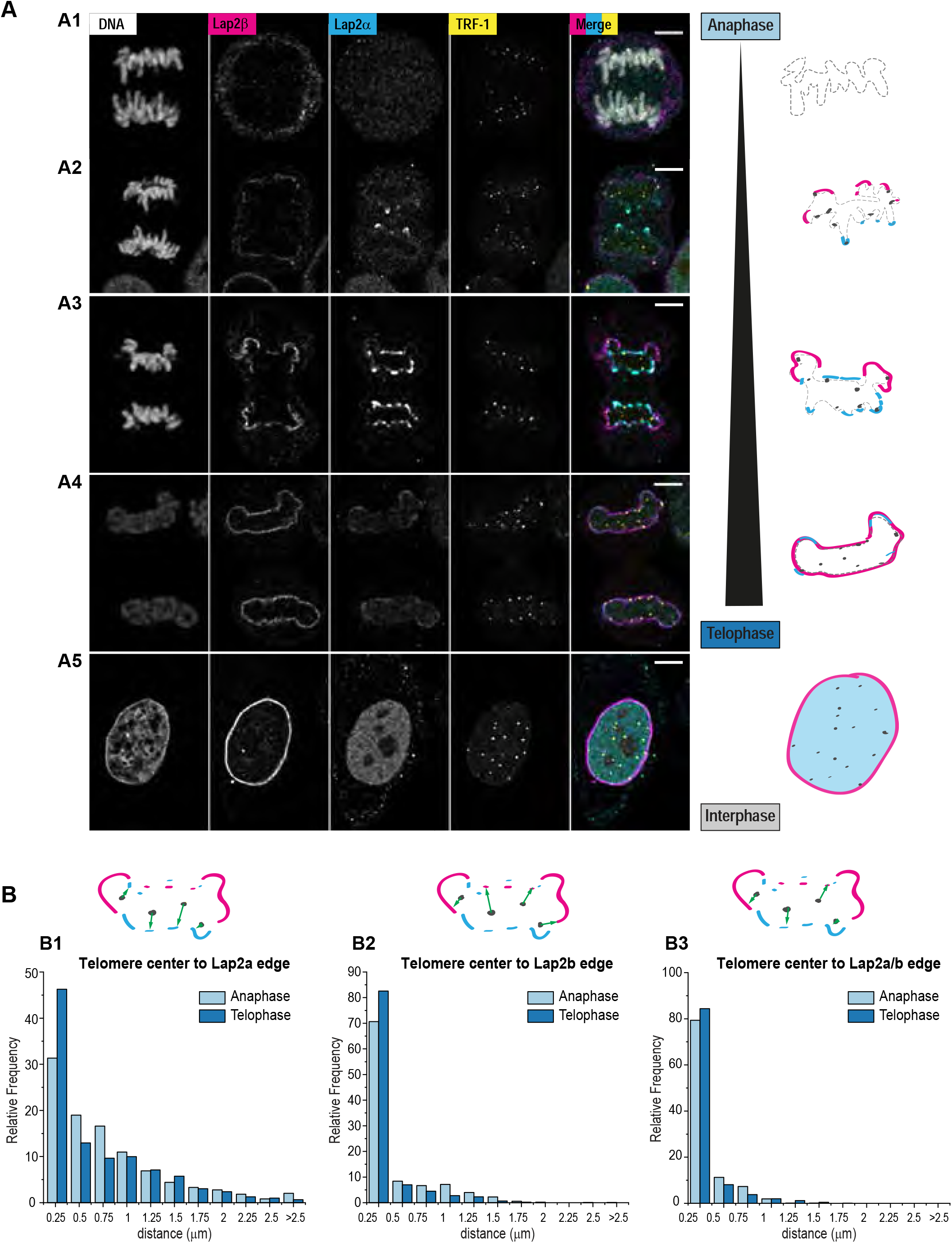
Lap2α and Lap2β are recruited to telomeres in early anaphase. A. Representative images and corresponding schematics illustrating the step-wise recruitment of Lap2α (*cyan*) and Lap2β (*magenta*) at telomeres (*TRF-1* – *yellow*) and chromosomes (DNA – *grey*) from anaphase to interphase. After being first un-detectable at anaphase onset (panel A1), Lap2α enriches at chromosome ends at telomeres and Lap2β is weakly detected at peripheral DNA (panel A2) followed by stronger enrichment of Lap2α at the core and Lap2β extends at the periphery (panel A3) until in Telophase all the DNA is encapsulated by Lap2β and Lap2α decreases at the nuclear envelope (panel A4). In interphase, Lap2β remains enriched at the nuclear envelope and Lap2α appears primarily nucleoplasmic (panel A5). Scale bars: 5 μm (main).Histograms of relative frequencies (%) of telomeres’ average distances from the center of mass of the segmented signal to the edges of segmented signal from Lap2α (B1), Lap2β (B2) or a segmentation mask combining both stainings (B3: Lap2a + Lap2b) measured for deconvolved confocal images of cells in anaphase (typically with morphologies as in A3) and in telophase (morphologies as in A4). Error bar indicates standard error of the mean; N=8 cells in anaphase, N= 12 cells in telophase; number of segmented telomeres per cells vary and are not indicated.

### BAF and LAP2 proteins affect 3D genome organization

We next decided to evaluate the outcome of disturbing the initial recruitment of LAP2α/β and BAF to telomeres. For this purpose, we combined siRNA directed against LAP2 proteins and BAF, and assessed the consequences using different readouts. We confirmed the efficiency of the knockdown by western blotting using whole-cell protein extracts (Fig. 5A). BAF is known to restrict accessibility of nuclear membranes to the surface of chromatin, and its depletion results in a global decrease of the nuclei circularity, because of lobular nuclear protrusions (Samwer et al., 2017). We therefore measured the circularity score for siLAP2, siBAF, and siBAF-LAP2 cells. Each single siRNA impacted nuclear shape, with a cumulative effect of the co-depletion (Fig. 5B – circularity < 0.6). Then, we evaluated the consequences of LAP2 and BAF codepletion on chromatin organization with MadID. We followed the same process as for the targeted screen, except that we analyzed the enrichment of LAD-CFHR3 together with two additional well-described LADs, CYP2C19 and CDH12, and used two independent iLADs regions to calculate relative enrichments. Both BAF and LAP2α single mutants increased the LaminB 1 contact frequency of LAD-CFHR3 in the screen (Fig. 3D), which was further increased upon knockdown of both proteins (Fig. 5C). In contrast, LAD-CYP2C19 and CDH12 were found less enriched at the NE in the double mutant compared to controls. These results were not a consequence of changes in the methylation status of iLADs after siRNA depletion (Fig. S8A). The depletion of LAP2 and BAF decreased telomere-NE attachment, also with an additive effect compared to single knockdowns (Fig. 3D, 5C). While these results support the implication of BAF and LAP2 in genome organization, they only provide a snapshot of the contact frequencies of a large population of asynchronous cells. To further address this phenotype in single cells and at different phases of the cell cycle, we applied our 3D-SIM spatial modeling approach (Fig. 2). In general, morphometric descriptors such as the number of telomeres, their density and nuclear sphericity were comparable in siRNA cells compared to controls (Fig. S8B) and compared to the initial dataset (Fig. S1D). Only the size of early G1 nuclei was reduced in siRNA BAF and LAP2, a phenotype that was recovered in G1/S cells (Fig. S8B). Surprisingly, we did not observe a significant drop of the fraction of telomeres localized within 500 nm of the NE in early G1 (Fig. 5D-E, S8C). This could be explained by the multiple tethering mechanisms foreseen for human telomeres and supported by the results of our screen and previous work (Crabbe et al., 2012). In addition, while the LAP2 and BAF depletion reached at least 80% in both single and double siRNA, immunostaining on fixed cells highlighted remains of LAP2α signal in anaphase, which remarkably colocalized with telomeres (Fig. S7B). This LAP2α leftover found at chromosome tips of LAP2α-depleted cells could maintain a certain level of telomere tethering. We did observe, however, consequences of LAP2 and BAF loss in interphase cells. The proportion of telomeres permanently found in proximity of the NE in interphase was reduced after LAP2 and BAF co-depletion (Fig. 5D-F, S phase) and spatial statistical analysis showed reduced positive spatial interaction with the NE (Fig. S8C), supporting the decrease in telomere enrichment seen by MadID. Overall, these experiments demonstrate that BAF and LAP2 proteins are involved in setting 3D genome organization. Their loss affects telomere tethering to the NE and changes the organization of LADs regions.

**Fig. 5.**
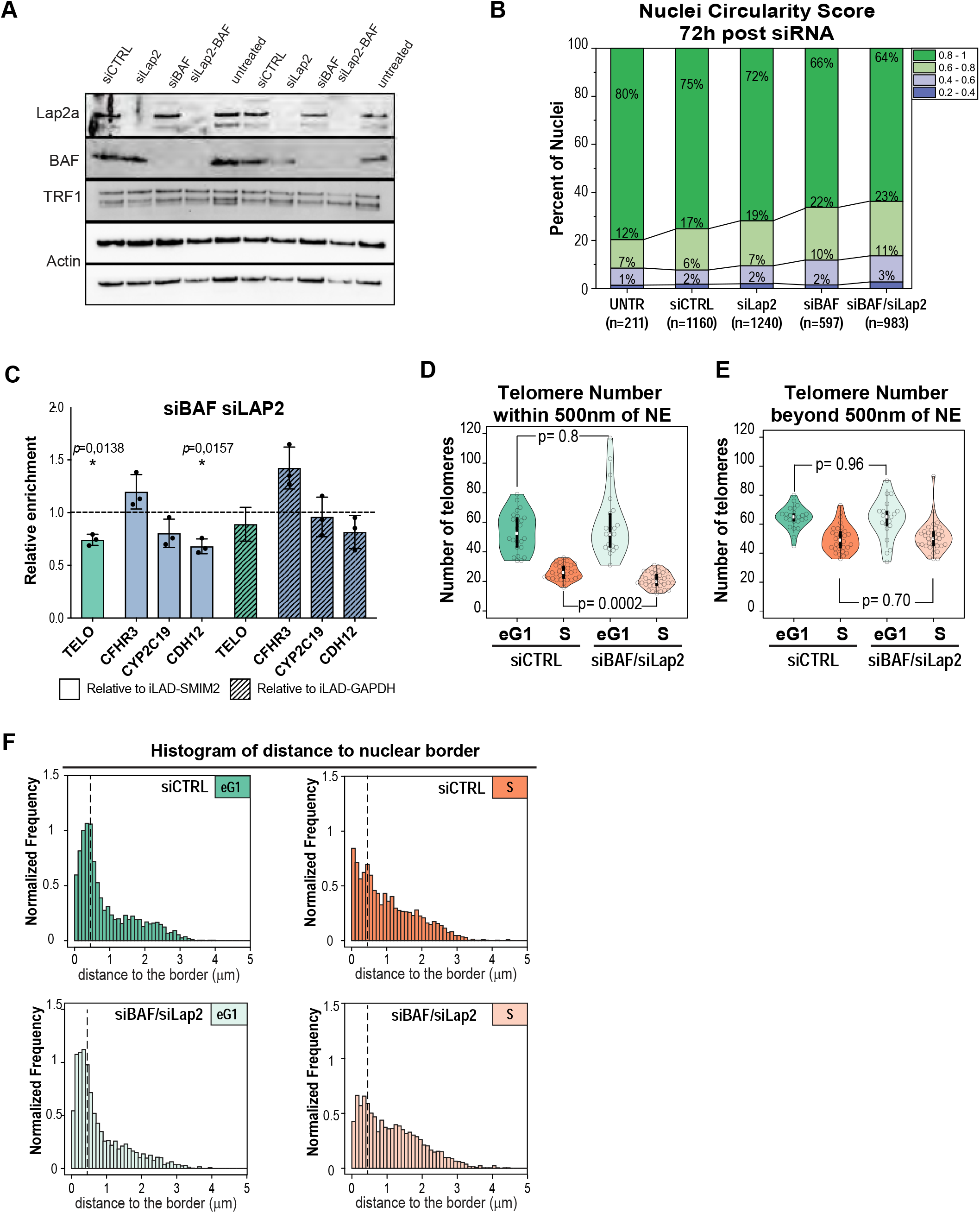
Depletion of Lap2α and Lap2β by siRNA affects telomere proximity to the nuclear envelope. A. Western blot of whole cell extracts showing the decreased protein levels of Lap2α or BAF in cells treated with siRNA against Lap2α (siLap2) or BAF (siBAF) or both (siLap2-BAF) compared to scrambled siRNA (siCTRL) or untreated cells. TRF1 is un-affected, actin serves as loading control. B. Changes in nuclear circularity 72h post-treatment, with an increase in abnormal shapes (circularity between 0.2-0.6) in siRNA treated cells compared to control. Number of analysed cells is indicated below from at least 2 independent experiments. C. Relative telomere, LAD-CFHR3, LAD-CYP2C19 and LAD-CDH12 enrichment calculated over iLAD-SMIM2 or iLAD-GAPDH and control condition (set at 1) in BAF- and LAP2-depleted cells. n=3. Mean ± SD is shown. D. and E. Effect of double siBAF/siLap2 on the total number of peripheral (D) and internal (E) telomeres in early G1 and S phases, with a significant decrease of periphal telomeres in doubledepleted siBAF/siLap2 cells in S phase. F. Distribution of the distance between each telomere and nuclear border in control (*Top*) and siBAF/siLap2 treated (*Bottom*) nuclei in early G1 and S phases indicating a decreased frequency of telomeres associated to the nuclear envelope (below dotted bar indicating 500 nm cut-off).

## Discussion

Genome organization is key to cellular identity, and structural nuclear landmarks need to be faithfully transmitted over cell generations in any given cell lineage. Rebuilding a functional G1 nucleus at the end of mitosis is therefore an essential process that requires the orchestration of numerous pathways, from NE reformation to re-establishment of chromosome territories and nuclear compartments. Here, we demonstrate that an interplay between NE proteins and telomeres impacts genome architecture, highlighting a new essential role for the ends of linear chromosomes.

### Telomeres and NE reformation

Our data indicate that telomeres are part of these discrete chromatin regions that attract a specific set of NE proteins in late anaphase, to initiate membrane formation. These interactions could represent the initial contacts between telomeric chromatin and the NE that perpetuate after mitotic exit, until early G1. Indeed, we estimated that ~80% of telomeres are located less than 250 nm from LAP2α/β anaphase patches at the onset of NE reformation (Fig. 4B), and that more than 50% of telomeres are localized within 500 nm of the nuclear border in early G1 cells (Fig 1C), which correspond to the estimated thickness of the nuclear lamina. What drives the initial recruitment of LAP2α on anaphase telomeres is unclear. An earlier study proposed that after NE breakdown, residual foci of LaminB1 were found throughout mitosis at major nuclear compartments, such as chromatin (Martin et al., 2010). These landmarks could be targeted during NE reassembly to impart spatial memory from one cell cycle to the next. Even though we were not able to detect these LaminB1 remnants in our cellular model, it would be of interest to see how telomeres organize with regards to the NE as they enter mitosis. A similar scenario was recently suggested for the heterochromatin mark H3K9me2, which acts as a 3D architectural mitotic guidepost, by orchestrating spatial repositioning of heterochromatin through mitosis (Poleshko et al., 2019a). This could suggest that the initial recruitment of NE proteins depends on the presence of specific heterochromatin marks at telomeres, such as H3K9me2/3, which are involved in perinuclear heterochromatin attachment to the nuclear lamina (Kind et al., 2013; Poleshko et al., 2013). While a mix of both heterochromatic and euchromatic marks characterizes the epigenetic status of telomeres (Cubiles et al., 2018), the presence of H3K9me2/3 has been observed at telomeric repeats in several human cellular models (Arnoult et al., 2012; Chow et al., 2018; Kychygina et al., 2021a; O’Sullivan et al., 2010). Notably, H3K9me3 can recruit heterochromatin protein 1 (HP1) (Arnoult et al., 2012), a major actor in tethering heterochromatin to the nuclear lamina (Poleshko et al., 2013). It was also demonstrated that H3K9me2 is retained through mitosis to reposition LADs at the nuclear periphery in daughter nuclei. During mitosis, H3K9me2 is hidden by the phosphorylation of H3 serine 10 (H3S10P), which drives the release of these regions from the nuclear periphery (Poleshko et al., 2019b). Genetic tools could be used to assess the role of these marks on telomere tethering, such as the forced enrichment of HP1α at telomeres, already known to induce telomere structure irregularity (Chow et al., 2018).

### 3D telomere distribution is not random

Using 3D spatial statistical modeling, we found that telomeres obey a complex spatial organization in the nuclear volume and showed that beyond a radial organization, they follow a globally repulsive, regular distribution associated in early G1 with mutual attraction in the short distance range. While repulsion was maintained throughout the cell cycle, local attraction vanished beyond early G1 and the radial patterning dynamically evolved throughout the cell cycle, suggesting independent determinants for these spatial organization features.

The physical location of telomeres at the extremities of the p and q-arms of each chromosome could potentially account for the spatial repulsion between telomeres, though it can be expected that the corresponding genomic distance would at least partially be compensated by the Rabl-like organization we noticed in early G1. Since chromosomes are organized into distinct, non-overlapping chromosome territories in interphase nuclei (Cremer and Cremer, 2010), an alternative is that repulsion would result from the physical separation of telomeres located on distinct chromosomes, in particular if telomeres are non-uniformly distributed relatively to their respective chromosome territories. The localization of telomeres at the periphery of their chromosome territories (Sehgal et al., 2016), the polar organization of chromosome territories with opposite centromeric and telomeric ends (Amrichová et al., 2003b), and non-random orientations of telomeric ends (Schmälter et al., 2014) are consistent with a non-random positioning of telomeres relatively to their respective territories. The maintenance of long-range repulsion we observed during the whole cell cycle as opposed to the vanishing short-range attraction after early G1 is consistent with previous reports of absence of large-scale reorganization of chromosome territories beyond G1 but of more dynamic patterns at the sub-chromosomal scale (Essers et al., 2005; Müller et al., 2010; Sehgal et al., 2016; Walter et al., 2003a).

Thanks to the resolution offered by 3D-SIM and the large number of characterized nuclei, we were able to achieve a high-resolution analysis of the distance between telomeres and nuclear periphery. This analysis revealed a bimodal distribution showing the existence throughout the cell cycle of two sub-populations of telomeres, one peripheral and one more internal. Chromosome territories in interphase nuclei distribute radially according to size, gene density, or transcriptional activity (Bolzer et al., 2005; Cremer et al., 2001; Mayer et al., 2005). Given the non-random localization of telomeres with respect to their territory, concentric layers of chromosomes could thus induce different modes in telomere distance to nuclear border. One striking result of our spatial analyses was to reveal that this interaction between telomeres and the nuclear border is dynamic, from a preferentially peripheral location in early eG1 to a preferentially internal one in G2. This is consistent with a previous 2D analysis study (Crabbe et al 2012) and recent results showing interactions between distal LADs and nuclear periphery decreased during cell cycle (van Schaik et al 2020).

### Inheritance of chromosome positions through mitosis

While it is well admitted that chromosomes in mammalian cells form territories, i.e. mutually exclusive globular volumes within the nuclear volume, their precise transmission from mother to daughter cells is still controversial (Bickmore and Chubb, 2003; Walter et al., 2003b; Williams and Fisher, 2003). Earlier work using elegant 4D imaging suggested that chromosome positions are transmitted through mitosis (Gerlich et al., 2003). However, it was also reported that the position of chromosome domains is not precisely passed on through mitosis but is rather plastic and redefined during early G1 (Thomson et al., 2004). Since the nuclear lamina serves as an anchor to organize chromatin, studies addressing single-cell genome-lamina interactions based on the DamID technology have been instrumental to shed light on these mechanisms (Kind et al., 2015). Collectively, LADs from all chromosomes cover ~30-40% of the mammalian genome (Guelen et al., 2008; Peric-Hupkes et al., 2010), but only ~30% of LADs are found at the periphery in each individual cell nucleus (Kind et al., 2013). These results suggest that LAD interactions with the NE are dynamic, with some LADs making permanent contacts, and other being more variable. They also suggest that even if nuclear organization is not entirely conserved and transmitted through cell divisions, a certain level of organization is preserved. Essential events occurring during mitosis, first during chromatin condensation in prophase to expose or hide certain chromatin loci, then in late anaphase to reform the lamina at accessible discrete sites, surely set 3D organization for the next interphase. Here, we show that telomeres, and probably extended regions at the ends of linear chromosomes, have this property. One interesting hypothesis is that association of telomeres with forming nuclear lamina in telophase could pull on the corresponding chromosome during nuclear expansion in early G1 and favor its peripheral position. In agreement with this view, genomic repositioning assays using ectopic LAD- or non-LAD-derived sequences expressed in cells clearly showed that mitosis is required for the interaction of a LAD sequence and the lamina (Zullo et al., 2012). While the ectopic LADs were localized at the periphery of the condensed chromosomal regions in early stages of mitosis, non-LAD sequences were found within the chromatin mass. This organization of condensed chromatin might favor accessibility in later stages, when the lamina is reforming around chromatin. Indeed, at the end of anaphase, a striking colocalization of LaminB1 with the LAD-derived sequence was observed (Zullo et al., 2012), similar to what we observe between telomeres and LAP2α.

### Molecular mechanisms driving telomere-NE attachment

The targeted screen we present here suggests that several pathways are at play to tether telomeres to the NE. This is coherent with the fact that mechanisms by which peripheral heterochromatin is tethered to the lamina are redundant (Olins et al., 2010; Poleshko et al., 2013; Solovei et al., 2013), with at least three mechanisms that rely on adaptor complexes that include an inner nuclear membrane protein such as BAF, LAP2α, or LaminB Receptor. DamID has again been instrumental to dig into this question further. When applied with the methyltransferase fused to LaminB1, LaminB2, LaminA or BAF, highly similar maps of LADs were obtained. Additional experiments modulating the levels of LaminA and BAF revealed that these tethering proteins compete for LAD binding (Kind and Steensel, 2014). In contrast, we did not observe an increased frequency of contacts between telomeres and LaminB1 after depletion of LaminA or BAF, as if alternative mechanisms could not compensate each other. Whether telomeres from specific chromosome arms, known to carry distinctive subtelomeric regions, are tethered by different pathways remains to be addressed. While centromeres kept their peripheral localization, the enrichment of telomeres at the nuclear rim was largely lost as soon as cells reached interphase, once nuclei regained their mature size and shape. A similar behavior was recently described for LADs using pA-DamID, a modified version of the CUT&TAG method (Kaya-Okur et al., 2019) combined with DamID (Schaik et al., 2020). Applied to synchronized cells, the authors could analyze the cell-cycle dynamics of lamina-associated DNA sequences. They proposed that distal chromosome regions up to ~25 Mb from telomeres are in contact with the nuclear lamina within the first hours after mitosis, before they gradually move away from the nuclear edge.

A small subset of telomeres retained their NE-anchoring throughout interphase. Using MadID combined with whole genome sequencing performed on asynchronous cells, we could previously highlight a preferential enrichment of some subtelomeric regions close to LaminB1 (Sobecki et al., 2018), corresponding to middle- or late-replicating telomeres (Arnoult et al., 2010). It is likely that the mechanisms of telomere tethering differ whether they occur during postmitotic assembly, or throughout interphase. This could explain why the proximity of telomeres to the reforming NE in telophase persisted after LAP2 proteins and BAF depletion, but we still observed a change in telomere distribution in interphase.

## Supplemental Figure Legends

**Fig. S1. Sample preparation, imaging, and quantification using 3D-SIM.**

A. Timeline showing main steps of sample preparation and fixation after cell synchronization using double-thymidine block (see Materials & Methods): S cells were collected at the end of the 2^nd^ thymindine block (at the G1/S boundary); G2 cells were collected 6h after release and early G1 (eG1) cells were collected 8.5h after release.

B. Representative images of nuclei markers imaged using 3D-SIM in sample nuclei at different phases of the cell cycle. SUN-1 staining is used to distinguish cells in early G1 specifically, which exhibit different morphologies: one “telophase like” with a more compact nucleus, and a “round” one. Merge panel contains LaminA/C (*magenta*), TRF1 (*green*) and SUN-1 (*yellow*) channels. Scale bar: 5 μm.

C. 3D-SIM raw image quality was assessed using SIMCheck (Ball et al., 2015b) and the modulation contrast value (MCNR) of the raw image was plotted for each channel imaged: DNA (DAPI), TRF1 (Atto 488), Lamin A/C (Dylight550) or SUN-1 (Alexa647). Each dot corresponds to one image, black bar indicates median value, dotted line indicates threshold for acceptance. N=61 cells (from the siRNA-treated dataset)

D. Quantitative 3D image analysis: nucleus size (D1) and shape (D2), total number (D3) and density (D5) of telomeres, number of peripheral (D7) and internal (D8) telomeres, telomere size (D4) and shape (D6) during cell cycle. Median values are indicated on graph. (N = 54, 43 and 39 for early G1, S and G2 phases, respectively).

E. Same as D for nuclei with labeled centromeres. (N = 35, 36 and 38 for early G1, S and G2 phases, respectively)

**Fig. S2. Statistical spatial analysis of peripheral and polar organization of telomeres and centromeres during cell cycle.**

**A.** Distance measurements used in spatial statistical analyses. Each telomere or centromere pattern (observed or simulated according to the random or the orbital model) was quantitatively characterized using distribution functions of distance measurements (blue lines). The measurements were: distances to nuclear border (function B); distances from nuclear center (function C); distances between arbitrary positions within the nuclear space and their nearest object (function F); distances to nearest neighbor (function G); distances to all other objects (function H).

**B.** Individual (*Top*) and population (*Bottom*) analysis of spatial interaction between telomeres and nuclear border. *Top*: observed cumulative distribution function (CDF) of the distance between each telomere and the nuclear border (function B; *Blue curve*) in a representative (median) nucleus at different phases of the cell cycle. *Orange*: average CDF computed over 99 patterns simulated according to a completely random model of telomere distribution. The 95% confidence envelope around the average was computed using another 99 simulated random patterns. *Bottom*: population histograms of the SDI computed using function B and the associated computed p-value (Kolmogorov-Smirnov test of uniformity).

**C.** Same as A for the analysis of the spatial interaction between telomeres and the center of the nucleus. This interaction was analysed based on the CDF of the distance between each telomere and the nuclear centroid (function C).

**D.** Same as A for centromeres.

**E.** Same as B for centromeres.

**Fig. S3. Polarity analysis of telomere and centromere nuclear organization across the cell cycle**

A. 3D views of segmented sample nuclei (*Color surfaces*) and their telomeres (*Red spots*) at different phases of the cell cycle. Nuclear surfaces are displayed opaque in front and rear views and are transparent in side views. The minor axis is orthogonal to the plane of view in front and rear views.

B. Same as E for centromeres.

C. Computation of the polarity index along the minor axis of the nucleus. Positions are projected along the minor axis and the proportions of projections located above and below the center of the nucleus are computed. The polarity index is the largest of these two proportions.

D. Cell-cycle distribution of the polarity index along the minor (*Left*), major (*Middle*) and the intermediate (*Right*) nucleus axis for telomeres. Kruskal-Wallis rank sum test (KW-test) and post-hoc comparison tests (Wilcoxon test with Benjamini-Hochberg correction for multiple testing; W-test) were performed as indicated.

E. Same as D for polarity index along the minor axis for centromeres.

**Fig. S4. Statistical spatial analysis of peripheral organization of telomeres during cell cycle: difference between observed and predicted distribution functions**

A. Example of difference between the observed function G (Blue) in an individual nucleus at early G1 and the corresponding function G_0_ (Orange) according to a completely random model of telomere distribution. The red curve shows the difference between G and G_0_.

B. Differences between observed individual functions G and associated model-predicted average functions G. Each color corresponds to a different nucleus. Predictions were computed using a completely random model of telomere distribution. The intervals were G-G0>0 (G-G0<0) highlight an excess (deficit) of measured distances in observed patterns as compared with model predictions. For example, G-G0>0 in the short distance range points to an attractive trend.

C. and D. Same as (A) with functions H and F.

**Fig. S5. M.EcoGII-driven DNA methylation is targeted and detected specifically.**

A. Immunofluorescence staining LaminA/C and v5-tagged M.EcoGII in M.EcoGII or M-LB1 expressing cells, in absence or presence of Shield1 (1 μM, 24h). Scale bar: 10 μm.

B. DNA and protein immunofluorescence staining methylated N6-adenosine (m6A) and LaminA/C in M.EcoGII or M-LB1 expressing cells, in absence or presence of Shield1 (1 μM, 24h). Scale bar: 10 μm.

C. Immunoblot to detect methylated N6-adenosine (m6A) in genomic DNA extracted from M.EcoGII or M-LB1 expressing cells, in absence or presence of Shield1 (1 μM, 24h).

D. Mean Ct values of three technical replicates obtained upon telomere, LAD-CFHR3 or iLAD-SMIM2 amplification by qPCR in control condition. n=16. Mean ± SD is shown.

E. Relative telomere and LAD-CFHR3 enrichment calculated over iLAD-SMIM2 in control condition. n=16. Mean ± SD is shown.

F. Immunofluorescence staining LaminA/C and v5-tagged M.EcoGII in control, FARα- or FARβ-depleted M.EcoGII or M-LB1 expressing cells treated with 1 μM Shield1 for 24h. Scale bar: 10 μm.

**Fig. S6. Depletion of proteins from the screen**

S6.1. Western blots of total protein extract from M.EcoGII (M) or M.EcoGII-LaminB1 (MB) expressing cells transfected with the indicated siRNA. Actin was used as a loading control. The estimated percentage of depletion is indicated.

S6.2. Relative mRNA expression of the indicated targets from M.EcoGII (M) or M.EcoGII-LaminB1 (MB) expressing cells transfected with the indicated siRNA. Relative mRNA expression was calculated using different references: Tata Binding Protein (TBP), beta 2 microglobulin (β2M), Ribosomal Protein L (RPL), or Actine (ACT).

**Fig. S7. Recruitment of Lap2α and Lap2β at telomeres in anaphase**

A. Top panel: orthogonal views of deconvolved confocal images at different cell sections (top, middle, bottom) showing early recruitment of Lap2α (cyan) or Lap2β (*magenta*) proteins preferentially at or around telomeres (*yellow*). DNA in shown in *grey*. Lines indicate planes for orthogonal views in both main and inset.

Bottom panel: section of a deconvolved anaphase cell, highlighting Lap2α (cyan) recruitment around a telomere (see inset), with Lap2β (*magenta*) exhibiting a slightly different pattern.

B. Decreased signal of Lap2α (cyan) or Lap2β (*magenta*) proteins after siRNA depletion, but remnants of Lap2α are still preferentially recruited around telomeres in early anaphase.

Scale bars: 5μm in main, 1μm in inset.

**Fig. S8. BAF and LAP2 depletion affect 3D genome organization**

A. Telomere, LAD-CFHR3 and iLAD-SMIM2 enrichment in control (n=4), BAF (n=3), LAP2 (n=2) or BAF and LAP2 (n=3) depleted cells. Mean ± SD is shown.

B. Quantitative 3D image analysis of siBAF/siLap2 treated cells in early G1 and S phase (see Methods section): nucleus size (B1) and shape (B2), total number (B3) and density (B4) of telomeres. Statistically significant differences and corresponding p-values are indicated on graph.

C. Analysis of spatial interactions between telomeres and nuclear border (*C1*) or the nuclear center (*C2*) and between telomeres (*C3-C5*) using comparisons to the random model of telomere organization for control or Lap2/BAF siRNA treatments. *C1*: distribution SDI computed using the CDF of the distance between each telomere and nuclear border, or *C2* nuclear center. *C3-C5:* same as (*C1*) for SDIs computed based on the cumulative distribution function of the distance between each telomere and its closest neughbor (*C3*), of the distance between each telomere and any other telomere (*C4*), and of the distance between arbitrary nuclear positions and their closest telomeres (*C5*). (p: p-value of Kolmogorov-Smirnov test of uniformity).

## Materials & Methods

### Detailed Step-By-Step Protocol

#### Cell culture and treatments

HeLa 1.2.11 cells (Crabbe et al., 2012) were grown in Glutamax-DMEM (GIBCO) supplemented with 10% (v/v) fetal bovine serum (GIBCO), 1% (v/v) non-essential amino acids (GIBCO) and 1% (v/v) sodium pyruvate (GIBCO), at 7.5% CO_2_ and 5% O_2_. For cell synchronization, cells were treated with 2 mM thymidine (Sigma) for 16h, followed by 3x washes with pre-warmed PBS (Sigma), and released into growth medium for 8h. A second 2mM thymdine treatment was performed for 16h, followed by washes and release as above, and G1/S samples were collected 0h, G2 cells 6h and early G1 cells 8.5h after release. Expression of pRetroX-PTuner vectors carrying M.EcoGII, M-LB1 or M-TRF1 was induced by addition of 1 μM Shield-1 (Aobious) for 24 hours. Protein depletions were carried out by transfection of 5 μM short interfering RNA (siRNA) for 72h using DharmaFECT transfection reagent (Horizon). For co-depletions (3D-SIM samples), the concentrations of siRNA used were: 5 μM siCTRL (“siCTRL”), 2.5 μM siCTRL + 2.5 μM siLap2 (“siLap2”), 2.5 μM siCTRL + 2.5 μM siBAF (“siBAF”), 2.5 μM siBAF + 2.5 μM siLap2 (“siBAF/siLap2”). Samples were collected at indicated timepoints and processed further (protein extraction, RNA extraction, immunofluorescence).

#### MadID

MadID was essentially performed as described (Sobecki et al., 2018). Briefly, genomic DNA was isolated using the Blood & Cell Culture DNA Midi kit (QIAGEN) according to the manufacturers’ recommendations. Additionally, an RNase treatment was performed for 1 hour at 37°C: 200 μg/mL RNaseA (Sigma), and 2.5 U/mL RNaseA and 100 U/mL RNaseT1 (RNase cocktail, Ambion).

10 μg of genomic DNA were sonicated during 40 cycles (30 s ON/60 s OFF, low intensity) into 200-400 bp fragments using a Bioruptor Plus sonicator (Diagenode). Sonication efficiency was verified by electrophoresis in a 1.5% agarose gel. 3% of sample was taken as an input. After denaturation for 10 min at 95°C and incubation on ice for 10 min, samples were supplemented with 10x m6A-IP buffer (100 mM Na-Phosphate buffer, pH 7.0; 3 M NaCl; 0.5% Triton X-100), 2.5 μg m6A antibody (Synaptic Systems) and rotated overnight at 4°C. Next, 20 μL of protein A Dynabeads (Thermo Fisher Scientific), pre-blocked for 1 hour in 0.5% BSA and 0.1% Tween-20 in PBS, were added and samples rotated at 4°C for 3 hours. Beads were then washed 4 times in 1 mL 1x m6A-IP buffer. Beads and input samples were resuspended in 100 μL digestion buffer (50 mM Tris, pH 8.0; 10 mM EDTA; 0.5% SDS) containing 300 μg/mL proteinase K and incubated for 3 hours at 50°C while shaking. DNA was purified with the QIAquick PCR Purification kit (QIAGEN) and eluted in 50 μL.

#### Quantitative PCR

Samples obtained after MadID were diluted 1:10 before performing qPCR with the primers listed in Supp. Table. For LADs and interLADs regions, 3 μL of sample were used in a reaction mix containing 1 μL 10 μM of each primer (0.5 μM final) and 5 μL 2x Light Cycler SYBR Green LightCycler^®^ 480 SYBR Green I Master (Roche). For telomere amplification, the reaction mix was the same except that 0.2 μM of primer TelG and 0.7 μM of primer TelC were used. Amplification cycle for LADs and interLADs regions was: 15 min at 95°C followed by 40 cycle of 15 s at 95°C and 1 min at 60°C. Amplification cycle for telomeres was: 2 min at 50°C followed by 40 cycles of 15 min at 95°C, 15 s at 94°C and 1 min at 54, and an extension step of 30 s at 72°C.

Standard curves with either input or telomeric DNA were included in every qPCR to ensure amplification efficiency close to 100%. Telomeric DNA consisted of 800 bp of telomeric TTAGGG repeats purified from pSP73.Sty11 plasmid, a gift from Titia de Lange (The Rockefeller University, USA), upon EcoRI digestion. Enrichment were calculated using the △△Ct method. The first △Ct corresponds to the difference between the sequence of interest (LAD-CFHR3 or telomeres) and the iLAD-SMIM2 that is deprived of methylation in M-LB1 cells. The △△Ct was then calculated using M.EcoGII samples as a control average.

#### RT-qPCR

Messenger RNA (mRNA) extraction and cDNA synthesis were performed using the μMACS One-step cDNA kit (Milteny) according to manufacturers’ recommendations. RT-qPCR reaction mix was performed in 0.5 μM FW primer, 0.5 μM RV primer and 1x Light Cycler SYBR Green LightCycler^®^ 480 SYBR Green I Master (Roche). Amplification cycle was: 10 min at 50°C and 5 min at 95°C followed by 40 cycles of 10 s at 95°C and 30 s at 60°C. Standard curves were performed to verify that amplification efficiency was close to 100%.

#### Western blotting

Whole protein extracts were obtained by lysing cell pellets in 1x laemmli buffer (10% glycerol, 2% SDS, 63 mM Tris-HCl pH 6.8) followed by 2 cycles of sonication of 15 s. 30 to 35 μg protein in LDS lysis buffer NuPAGE (Thermo Fisher Scientific) were then heated at 95°C for 10 min and resolved on pre-cast 4-12% SDS-PAGE gradient gels (Invitrogen). Transfer of proteins from SDS-PAGE gels to nitrocellulose membranes was performed by dry transfer using the iBlot™ 2 Gel Transfer Device (Thermo Fisher Scientific) following the manufacturers’ recommendations. Primary antibodies were incubated for 2 hours at RT or overnight at 4°C, followed by 1-hour secondary antibody incubation. Membranes were overlaid with western blotting substrate for 5 min (Clarity™, BioRad) before visualization with a ChemiDoc™ Imaging System (BioRad).

#### m6A immunodot blot

Dot blot of genomic DNA was performed as described (Kychygina et al., 2021b) using the BioRad 96-well Bio-Dot^®^ apparatus. Positively charged Amersham Hybond-N+ membranes (GE Healthcare) and Whatman filter papers (GE Healthcare) preincubated with 2x SSC buffer were assembled onto the apparatus. Samples were denatured (98°C for 10 min followed by 10 min incubation on ice) and loaded on the membrane via vacuum blotting, followed by washing of the wells with cold 2x SSC. The membrane was denatured and neutralized sequentially by placing it on top of a Whatman filter paper (DNA face up) saturated with denaturing solution (1.5M NaCl, 0.5M NaOH) for 10 min at RT and neutralization solution (0.5 M Tris-HCl, pH 7.0, 3 M NaCl) for 10 min at RT. The membrane was crosslinked with UV at 120000 μJ/cm^2^ and blocked for 1 hour in 5% non-fat dry milk in 0.1% TBST (0.1% Tween-20 in 1x TBS, pH 7.4). Subsequently, m6A antibody (Synaptic Systems) was diluted to 1:2000 in 5% non-fat dry milk and 0.1% TBST, and incubated overnight at 4°C. Following three washes with 0.1% TBST, an HRP-conjugated secondary antibody was applied for 45 min at RT. Following three washes with 0.1% TBST, the chemiluminescence signal was visualized using the ChemiDoc Imaging System (BioRad). The intensity of the m6A signal was quantified using the ImageJ software.

#### Protein and DNA immunofluorescence

**For protein immunofluorescence (conventional microscopy),** cells were plated onto #1.5 coverslips and were fixed at given timepoints in 4% PFA in PBS for 10 minutes followed by permeabilization in 0.5% Triton 100-X in PBS. After three washes of 5 min in 1x PBS, cells were blocked for 30 min in PBG (1x PBS, 0.5% (w/v) BSA, 0.2% (w/v) cold water fish gelatin) followed by incubation with antibodies listed in Supp. Table (either overnight at 4°C or 2 hours at RT). After three washes of 5 min in 1x PBS and 45 min incubation with Alexa488/546/647-conjugated secondary antibodies (Invitrogen), coverslips were washed and mounted with Mowiol mounting medium (24% (w/v) glycerol, 9.6% (w/v) Mowiol 4–88, 0.1 M Tris-HCl pH 8.5, 2.5% (w/v) Dabco).

**For DNA immunofluorescence,** after fixation in 4% PFA and permeabilization in 0.5% Triton 100-X, cells were treated with 200 μg/mL RNaseA (Sigma), and 2.5 U/mL RNaseA and 100 U/mL RNaseT1 (RNase cocktail, Ambion) at 37°C for 1 hour. DNA was then denatured (1.5 M NaCl, 0.5 M NaOH) for 30 min at RT and neutralized (0.5 M Tris-HCl, pH 7.0, 3 M NaCl) twice for 5 min at RT. Samples were then washed three times with 1x PBS for 5 minutes at RT before proceeding with blocking and incubation of antibodies as described above.

**For protein immunofluorescence (3D-SIM imaging, deconvolved widefield & confocal),** cells were plated onto #1.5 High-Resolution coverslips (Carl Roth) pre-cleaned with a brief 100% ethanol wash and rinsed with distilled H20 prior to cell seeding. Immunofluorescence was performed according to the Kraus et al. protocol (Kraus et al., 2017) as follows: for fixation, cells were washed 2x with PBS before being transferred in a 6-well plate containing freshly prepared 4% formaldehyde (methanol-free; FA), preequilibrated at 37°C, and incubated for 10 min. FA was removed gently from one side, while simultaneously adding PBS + 0.02% Tween (PBSTw) on the other side for step-wise fixative exchange. Samples were washed twice more with PBSTw, quenched for 10min using freshly prepared 20 mM Glycine in H2O, followed by 2x PBSTw washes. After this stage, the samples were either put in PBS and kept at 4°C for further processing at a later time or were permeabilized for 10 min. in 0.5% Triton X-100. Samples were blocked for ~30 min in PBG before incubation with primary antibody or GFP-nanobody diluted in PBG, face-down on parafilm, in a humidified chamber (see antibody list below). After primary antibody incubation, samples were transferred in a 6-well plate and washed 4x with PBSTw, before being transferred in PBG with secondary antibodies at a dilution of 1:1000 for 45 min to 1h. After 4x PBSTw washes, samples were post-fixed with freshly prepared 4% PFA for 10 min and washed and quenched as described above. Hoechst was used as a DNA-counterstain at concentrations of 1-5 μm/mL for 5 min in H2O. A final wash was performed in H2O prior to mounting to remove any PBS remnant and avoid PBS crystal formation. The list of antibodies used can be found in **Suppl. Table** and the combination of primary & secondary antibodies used were: 1:100 GFP-nanobody conjugated with Atto488 (NanoTag) for TRF1-GFP, 1:1000 mouse anti-LaminAC (Santa Cruz) and horse anti-mouse Dylight549 or donkey anti-mouse Dylight 550, 1:350 rabbit anti-SUN-1 (Sigma) and/or rabbit anti-Cep152 (Sigma) and goat anti-rabbit Alexa-647; 1:1000 mouse anti-LaminAC (Santa Cruz) and horse anti-mouse Dylight488, 1:2000 human anti-CREST (Sigma) and donkey anti-human Dylight550, 1:350 rabbit anti-SUN-1 and goat anti-rabbit Alexa-647; 1:100 GFP-nanobody for TRF1-GFP, 1:250 mouse anti-Lap2a and anti-mouse Dylight549, 1:250 rabbit anti-Lap2b and anti-rabbit Alexa-647.

**3D-SIM mounting media**; given that the refractive index of the mounting medium plays a significant role in 3D-SIM image reconstruction quality, we developed a new mounting media based on a clearing agent - Sorbitol - that had been used for 3D-SIM imaging of DAPI-stained cells up to ~ 20 μm and shown to yield better modulation-to-contrast-to-noise ratio (MCNR) (Szczurek et al., 2018b). After empirical screening of additives and secondary antibodies, we used a formulation of 0.8x 70% wt/wt Sorbitol/PBS + 0.2x SlowFade Diamond (referred to as Sorb70SF) with a measured RI of 1.448 using Mettler Toledo 30PX refractometer at 23°C. Sorb70SF yielded bright and stable signal with DAPI, Dylight488 / GFP / Atto488, Dylight549 and Dylight 650/Alexa Fluor 647 (AF647), which we then used for the subsequent imaging. Sample mounting was performed as follows: coverslips were first pre-mounted in 35% wt/wt Sorbitol/PBS for 10 min, followed by 70% wt/wt Sorbitol/PBS for 5 min on parafilm. Coverslips were gently tapped onto tissue to remove excess sorbitol, mounted on a ~12-15 μL drop of Sorb70SF placed in the center of ethanol-cleaned unfrosted slides (Mänzel) and left to settle. Excess mountant was removed prior to sealing with nail polish, and the samples were kept at 4°C before imaging. For best results, samples were imaged within 1 week of preparation.

**Cell cycle staging for 3D-SIM siRNA-treated samples** was performed using SUN-1 speckles and the centrosomal marker Cep152, both stained with AF647 and imaged in the same channel. Early G1 cells were identified by the presence of SUN-1 peri-nuclear speckles (Fig. S1) and with 2 connected Cep152 rings (Park et al., 2014), while late S/ early G2 cells had SUN-1 signal only at the NE and exihibited 2 clear separated Cep152 rings connected to the NE, indicating the start of centrosome separation (Lawo et al., 2012; Piel et al., 2000).

##### Microscopy

**Widefield imaging** was performed on a OMX-SR (GE Healthcare/Cytiva) equipped with 405-, 488-, 568-, and 640-nm excitation lasers and emitted fluorescence was collected through a 60X 1.4 numerical aperture (NA) oil objective (Olympus) using an immersion oil of 1.514 refractive index (GE Healthcare) onto two pco.edge.4.1 scientific complementary metal-oxide-semiconductor (sCMOS) cameras (PCO), yielding a pixel size of 80 nm laterally. A dichroic was used to separate emitted fluorescence onto two separate lightpaths: emitted fluorescence was filtered using 431/31 nm or 528/48 nm emission filters for DAPI and GFP/Atto488/Dylight488 respectively on one camera, and 609/37nm or 685/40nm for Dylight 549 or AF647/Dylight650 on the second camera respectively. Z stacks (z-step of 262 nm) covering the 3D volume of the cells were acquired for multiple positions using the sequential mode “All Z then Channel”.

**3D-SIM images** were also acquired on the OMX-SR described above, with 3D stacks acquired over the whole cell volume with a *z* step of 125 nm and 15 raw images per plane (5 phases, 3 angles). Spherical aberrations were minimized by using immersion oil with refractive index (RI) of 1.516 - 1.520 for sample acquisitions. Images were acquired in sequential “All Z then Channel” mode: LaminA/C (Dylight549, 568 nm laser), TRF1 (FluoTag-Atto488, 488 nm laser), SUN-1 or Cep152 (AF647, 640 nm laser), DNA (Hoechst, 405 nm laser).

**Confocal images** for Figures 4 & S6 were acquired on a Leica TCS SP8 equipped with a 63x 1.4 NA oil objective, a pixel size of 58.5 nm and z steps of 298 nm. For Figure S4-A-B, images were acquired on a ZEISS Airyscan confocal microscope using a 63x oil objective with 1.4 NA and a pixel size of 155.8 nm.

For Figure S4-F, images were acquired on a ZEISS Airyscan confocal microscope using a 40x oil objective with 1.4 NA and a pixel size of 131.8 nm and z steps of 1 μm.

##### Image processing and analysis

**Image Deconvolution** of widefield (Fig. 5B) and confocal images (Figs. 4, S7) was performed using Huygens Essential (SVI). For widefield images, the following parameters were used: 50 max iterations, 40 SNR, 0.01 quality criterion and CMLE deconvolution mode. Chromatic shift correction was calculated in Huygens using a control cell stained only with Hoechst / DAPI and imaged sequentially with all lasers and fluorescing in all channels and deconvolved as above. The calculated chromatic shift correction was added to all deconvolved images. For confocal images deconvolution, the parameters were: 10 max iterations, 10 SNR, 0.05 quality criterion and CMLE deconvolution mode.

**3D-SIM Image processing and quantitative analysis:** 3D-SIM images reconstruction from raw data was performed with SoftWoRx v6.5.2 (GE Healthcare) using matched optical transfer functions (OTF) (Demmerle et al., 2017) recorded with 1.516 or 1.518 oil depending on sample thickness and Wiener filter settings set automatically, and channel-aligned using the SoftWorRx tool. Quality control on 3D-SIM images was done using different Fiji/ImageJ macros: first, images were thresholded in 16bit using a function within the SIMCheck plugin ((Ball et al., 2015b), *“1 SIMCheck_THR_DKO.ijm*”) then individual nuclei were cropped to ensure a single nucleus per image (“*2_CropNucleiBoundingBox.ijm”*), and finally the reconstruction quality was assessed via modulation contrast-to-noise ratio (MCNR) of SIMCheck, using a macro kindly provided by Lothar Schermelleh (“*3_SIMCheck_QC_EzMiron.ijm”*), both visually using the MCNR map on the image, and using the average value per image and channel. Only channels with and MCNR value >6 for the features of interest (LaminA/C, LaminB1, TRF1) were included in the analysis, and were plotted in Fig. S1 using “PlotsOfData” ((Postma and Goedhart, 2019), available at https://huygens.science.uva.nl/PlotsOfData/). The different channels of acquired 3D-SIM images were split under Fiji and stored in distinct TIFF files for further processing (“*SPLIT-Channels-into-Folders.ijm*”). All subsequent automated image processing procedures were implemented as batched pipelines using the bip software (http://free-d.versailles.inra.fr/html/bip.html).

The LaminA/C-Dylight549 channel was used to determine nuclear boundaries (BIP Pipeline *nucleus.pipeline*). Images were first resampled using linear interpolation to isotropic voxel size of 100 x 100 x 100 nm. Following Gaussian smoothing, a directional closing filter was applied to fill small gaps in the LaminA/C signal. Non-significant minima were removed by computing extended minima followed by minima imposition (Soille, 2004). The watershed transform was then applied on the resulting image (Vincent and Soille, 1991). Labeled regions touching the image borders were removed and the nucleus mask was obtained by selecting the largest remaining region. In a few cases, some artefacts remained (small holes or protrusions at the periphery), which were manually corrected under Fiji.

The TRF1-EGFP channel was used to extract telomeres (BIP Pipeline *telomeres.pipeline*). Following Gaussian smoothing, potentially remaining 3D-SIM-echo artefacts were removed using a specifically designed filter. With this operator, voxel values that were below some proportion of the maximal value of their neighbourhood were set to 0. A h-maxima operator was applied to remove non-significant intensity peaks. Automatic thresholding was then performed using an in-house operator specifically developed to extract constellations of objects of similar sizes, by selecting the threshold that minimized the coefficient of variation of object size. Following component labelling, telomeres not contained within the nucleus were removed by masking with the binary nuclear mask. The obtained image was used as a mask for the geodesic reconstruction of the unmasked image of labeled telomeres. This ensured that telomeres located close to the boundary of the nucleus were not truncated. In some cases, a few false positives remained in the images and were removed manually.

The CREST-Dylight550 channel was used to extract centromeres (BIP Pipeline *centromeres.pipeline*). Following Gaussian smoothing, a morphological size opening was used to remove small fluorescent spots. A first thresholding at a low threshold was applied to binarize the centromeric domains. A second thresholding at a high threshold was applied in parallel, followed by component labelling. The obtained labeled were then dilated within the mask obtained with the low threshold. This procedure ensured that centromeres in close vicinity were not aggregated under the same label. Masking and geodesic reconstruction were applied as for telomeres to select centromeres located within the nucleus of interest.

Morphological descriptors including volumes, centroid positions, and shape parameters (elongation, sphericity), were extracted from the segmented images. The nuclear boundary was represented as a triangular mesh extracted from the binary nuclear mask with the Marching Cubes algorithm (Lorensen and Cline, 1987). This surface was used for computing distances between object centers and nuclear border. Three-dimensional views of nuclear boundary and telomeric or centromeric positions were generated using the 3D viewer of the Free-D software (Andrey and Maurin, 2005); http://free-d.versailles.inra.fr).

##### Statistical spatial analysis

Telomeric and centromeric patterns were compared to distributions expected under theoretical models for the organizations of points (for telomeres) or real-sized objects (for centromeres) according to the methodology we recently developed (Arpòn et al., 2021b). Computer simulations were used to obtain these distibrutions. Nuclear boundary (triangular meshes) and telomere centroids were provided as input in these analyses. For centromeres, the measured individual centromere sizes were provided as additional input parameters and used to ensure that the simulated centromere positions were not leading to intersections between centromeres or with the nuclear envelope. Hence, the models took into account all the variables (nuclear shape and size, number and sizes of analyzed objects) that would otherwise bias the spatial analysis. In the completely random model, points or objects were distributed uniformly and independently (up to object intersections) within the nuclear space. In the orbital model, the relative positioning to the nuclear border was identical between observed and model-predicted patterns. The orbital model was indeed derived from the completely random model by keeping to its observed value the distance between each telomere and the nuclear border. The measured distances between telomeres and their closest points at the boundary of the nucleus were thus additional parameters provided as inputs to the orbital model. Comparisons between observed distributions of SDIs and expected uniform distributions were performed using the Kolmogorov-Smirnov test of distribution uniformity within the R software (R Core Team (2020). R: A language and environment for statistical computing. R Foundation for Statistical Computing, Vienna, Austria. URL https://www.R-project.org/.).

#### Polarity analysis

The three principal axes of each nucleus were computed from its binary mask and the coordinates of each telomere or centromere were expressed in the corresponding coordinate frame. The number N_i_^+^ (N_i_^-^) of telomeres or centromeres with positive (negative) *i*th coordinate was determined. The polarity index along the *i*th axis was computed as max (N_i_^+^,N_i_^-^)/(N_i_^+^+N_i_^-^).

**Nuclear Circularity analysis** was carried out using ImageJ/Fiji (Schindelin et al., 2012) and Cell Profiler (Stirling et al., 2021)), tools and pipelines can be found at: https://github.com/DeboraOlivier/Telomere3D/. Briefly, widefield imaged were projected along the Z axis according to the maximum intensity using a custom macro from the FILM facility, Imperial College London (https://www.imperial.ac.uk/medicine/facility-for-imaging-by-light-microscopy/software/fiji/). Nuclei and telomeres were segmented using a custom-made CellProfiler pipeline, and the circularity score was calculated for each nucleus after different treatments (untreated, siCTRL, siLap2, siLap2 + siBAF) using the *FormFactor* measurement in CellProfiler corresponding to 4*π*Area/Perimeter^2^ (Fig 5B), similar to (Appen et al., 2020) to quantify nuclear envelope reformation defects upon LEM2 siRNA. The results were then analysed using Origin2019.

**Confocal Image analysis (Figure 4):** Deconvolved confocal images where analyzed using BIP tool to segment LAP2A and LAP2B regions and telomeres and measure distances to LAP2 regions. An Euclidean distance map was computed on the inverted binary mask of LAP2 regions, providing the distance between any nucleus position and the closest LAP2-labeled position. This map was considered as an intensity image and intensity measurements were performed over segmented telomeres. For each telomere, the median distance value was retained as a measure of the distance between its center and the edge of the nucleus. The closest distance to LAP2 (A or B) was obtained by taking the minimum of the two distances to LAP2A and to LAP2B.

#### Data availability

The 3D-SIM images that have been generated in this work are publicly available from https://doi.org/10.57745/IQYEQS. Fiji & CellProfiler tools: https://github.com/DeboraOlivier/Telomere3D/ **Code availability**: All Fiji/ImageJ (Schindelin et al., 2012) & CellProfiler 4.0.7 (Stirling et al., 2021) image analysis tools can be found at: https://github.com/DeboraOlivier/Telomere3D/. Pipelines for the BIP software are available online on Recherche Data Gouv INRAE (https://doi.org/10.57745/0YF4AI).

All other relevant data supporting the key findings of this study are available within the article and its Supplementary Information files or from the corresponding authors upon reasonable request.

## Supplementary table. List of reagents used in this study

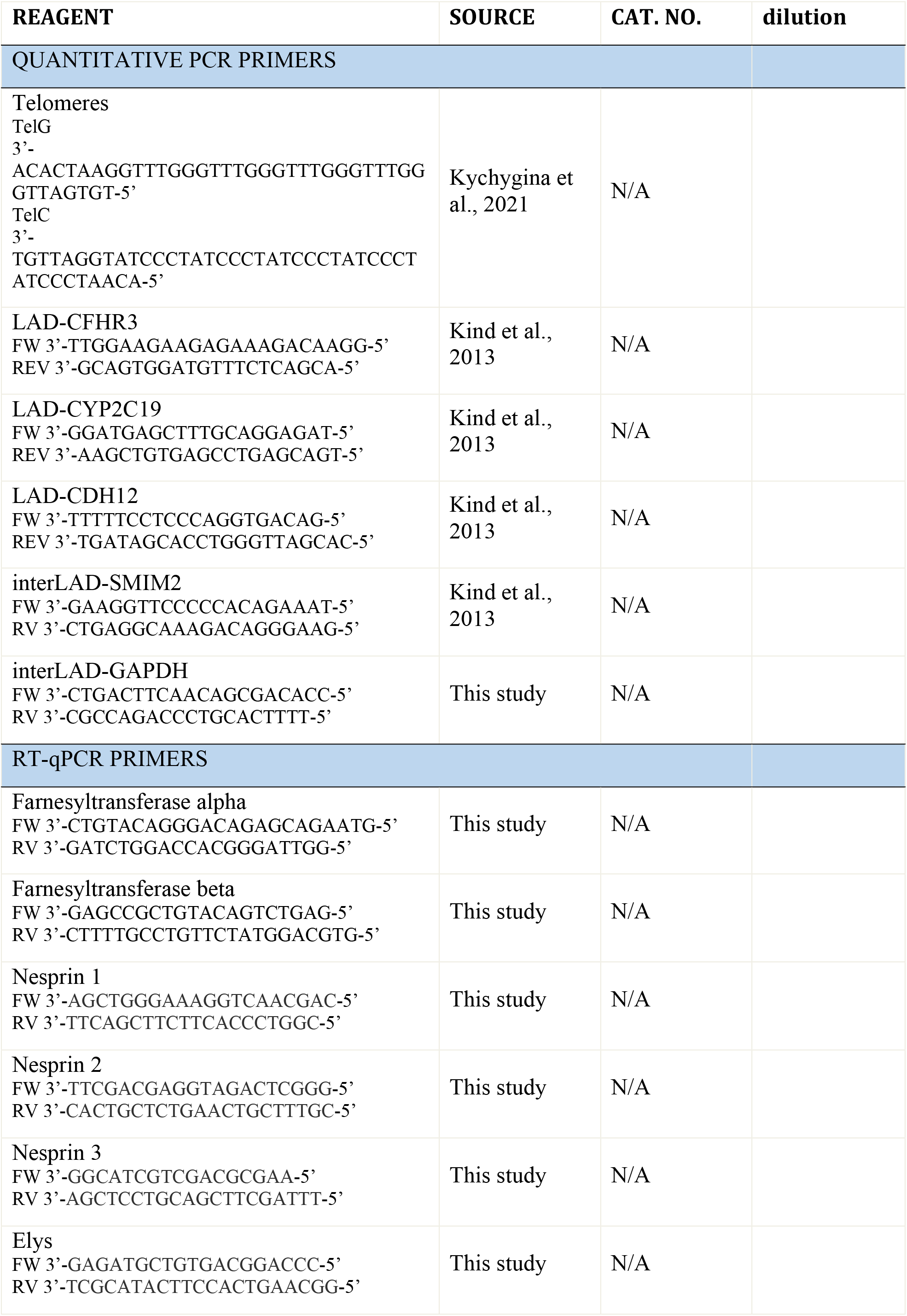
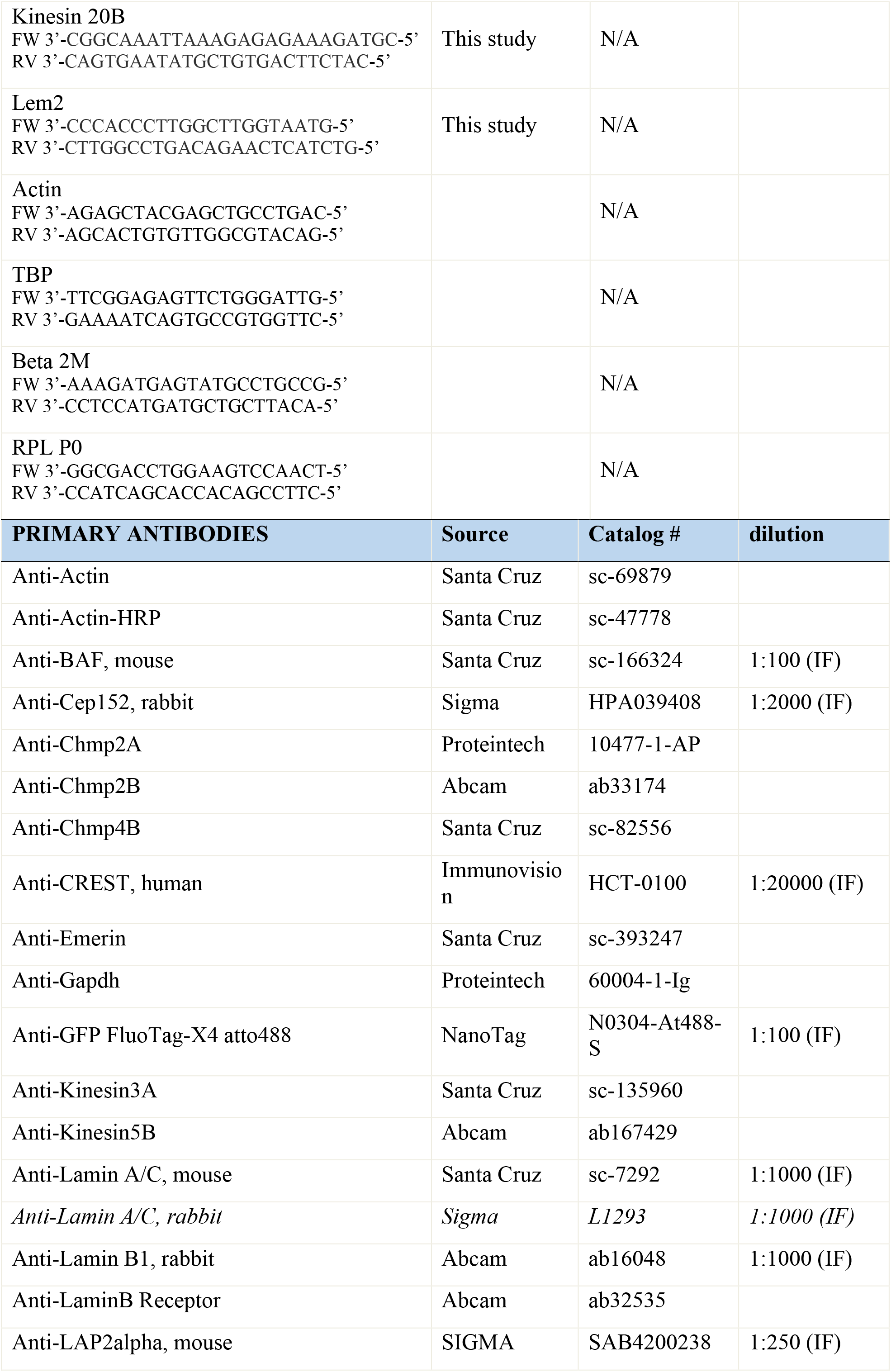
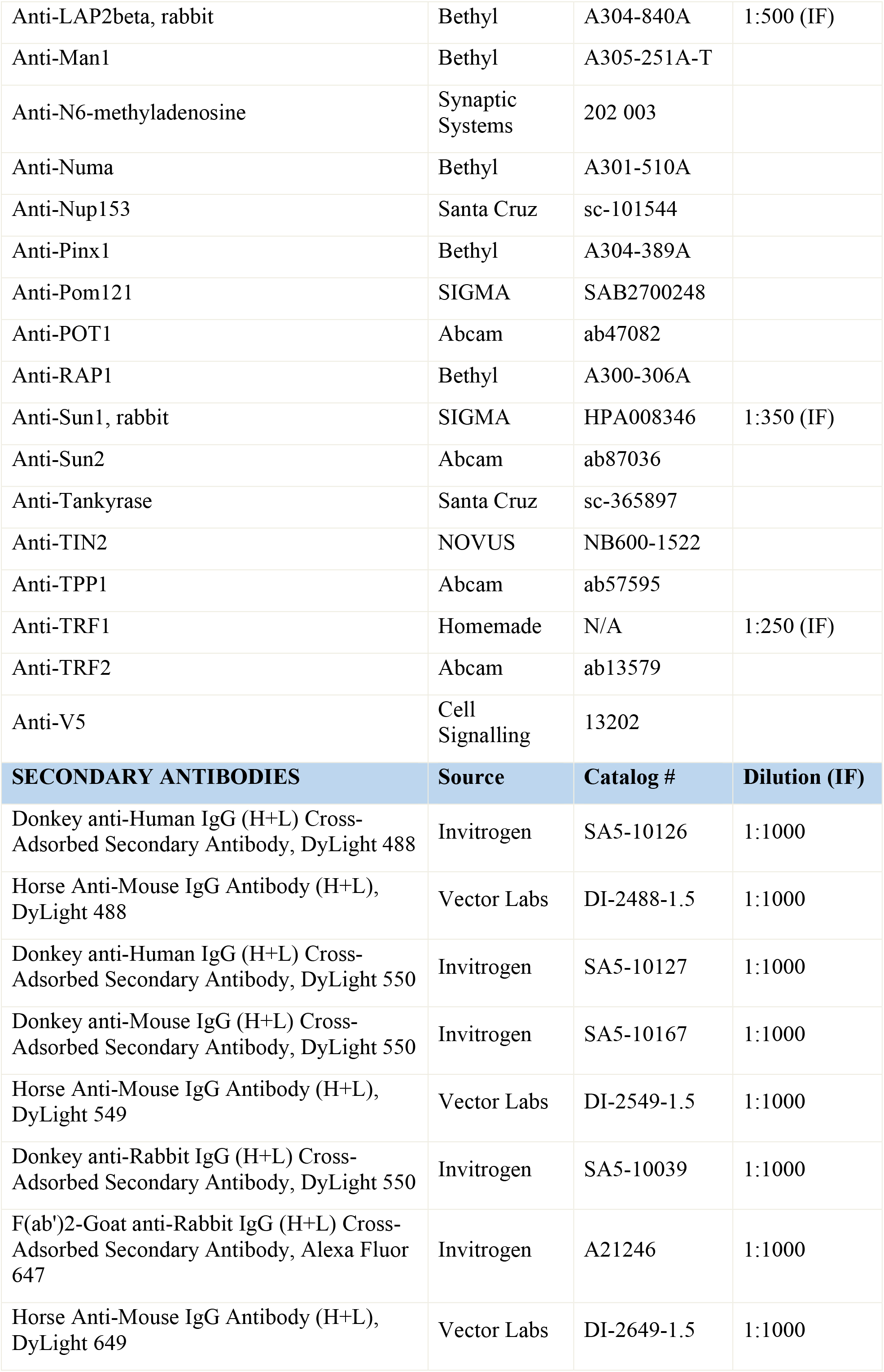
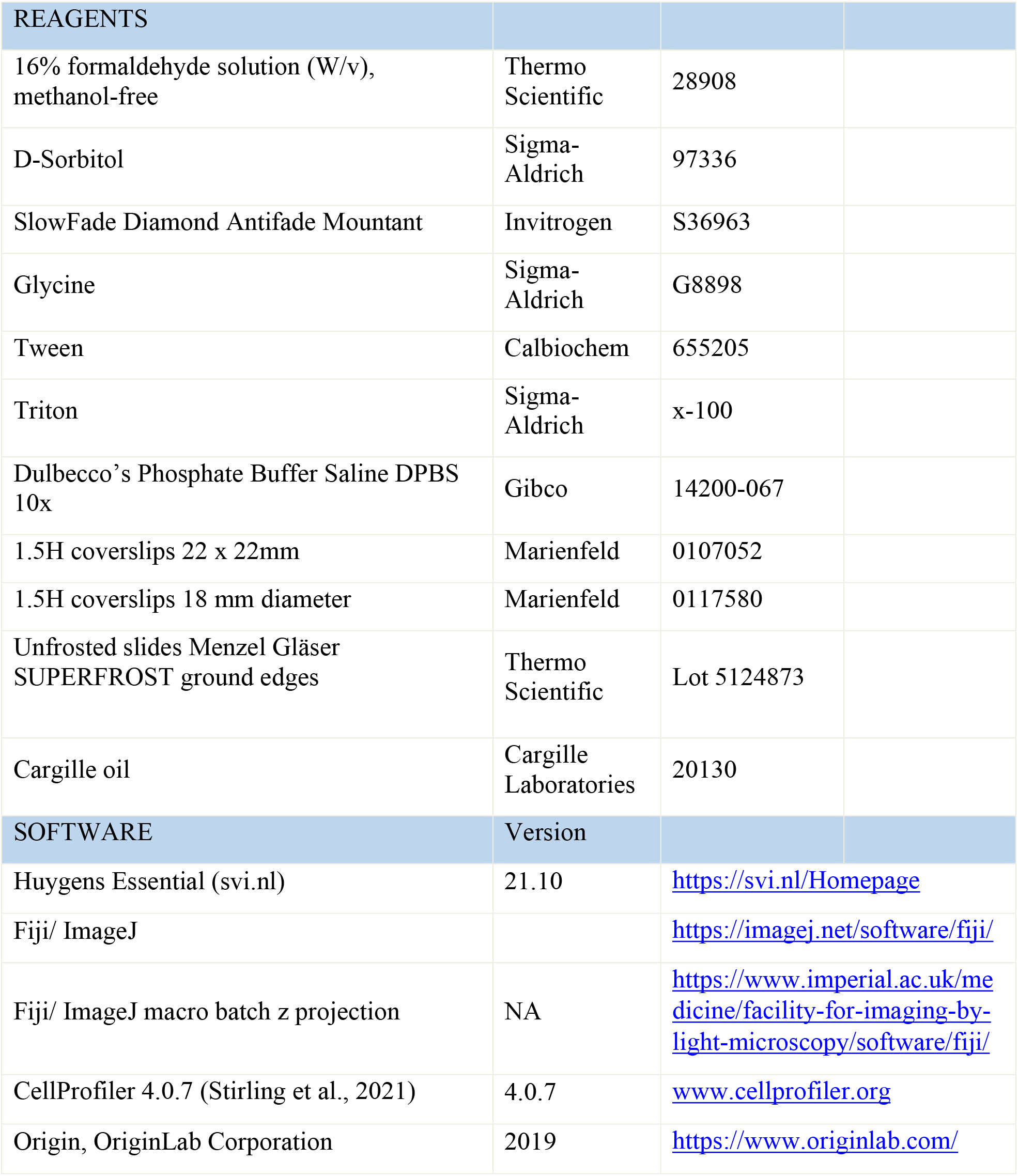

## Authors contribution (using CreDit Taxonomy)

Conceptualization, D.K., S.S. and L.C.; Methodology, D.K., P.A., P.M., S.S., D.U., L.C.; Investigation, D.K., N.O., S.S, D.U., L.C.; Formal Analysis, P.A.; Software, E.Bi. and P.A.; Writing – Original Draft, D.K., P.A., L.C.; Writing – Review & Editing, D.K., N.O., S.S, D.U., P.M., E.B., P.A. and L.C.; Funding Acquisition, L.C., P.A., E.B.; Resources, P.M. and E.B.; Supervision, L.C.

## Acknowledgements

We thank all members of the Crabbe laboratory for discussion, Anne Carpenter & Beth Cimini for training and help with CellProfiler tools, and Lothar Schermelleh for helpful discussions as well as providing the macro for SIMCheck. We thank Michal Sobecki for initially starting the subcloning of M.EcoGII-transduced cell lines. This work has benefited from the support of Morphoscope technological platforms at Ecole Polytechnique (Palaiseau). Work in the Crabbe lab was supported by the ATIP starting grant from CNRS in the framework of Plan Cancer 2014-2019 (to L.C.), the ANR Tremplin ERC teloHOOK (to L.C.), and the European Research Council grant teloHOOK/ERC grant agreement no. 714653 (to L.C.). EB, PM, DK by the ANR funding (ANR-11-EQPX-0029 Morphoscope2). The IJPB benefits from the support of Saclay Plant Sciences-SPS (ANR-17-EUR-0007). DK benefited from the COST Action (CA15124) - A Network of European BioImage Analysts to advance life science imaging (NEUBIAS) for a short-term scientific mission with Anne Carpenter (Broad Institute of Harvard and MIT).

## References

Adam N, Degelman E, Briggs S, Wazen R-M, Colarusso P, Riabowol K, Beattie T. 2019. Telomere analysis using 3D fluorescence microscopy suggests mammalian telomere clustering in hTERT-immortalized Hs68 fibroblasts. Communications Biology 2:451–12. doi:10.1038/s42003-019-0692-z

Adam SA, Butin-Israeli V, Cleland MM, Shimi T, Goldman RD. 2013. Disruption of lamin B1 and lamin B2 processing and localization by farnesyltransferase inhibitors. Nucleus 4:142–150. doi:10.4161/nucl.24089

Amrichová J, Lukásová E, Kozubek S, Kozubek M. 2003a. Nuclear and territorial topography of chromosome telomeres in human lymphocytes. Exp Cell Res 289:11–26. doi:10.1016/s0014-4827(03)00208-8

Amrichová J, Lukášová E, Kozubek S, Kozubek M. 2003b. Nuclear and territorial topography of chromosome telomeres in human lymphocytes. Exp Cell Res 289:11–26. doi:10.1016/s0014-4827(03)00208-8

Anderson DJ, Hetzer MW. 2008. The life cycle of the metazoan nuclear envelope. Current opinion in cell biology 20:386–392. doi:10.1016/j.ceb.2008.03.016

Anderson DJ, Vargas JD, Hsiao JP, Hetzer MW. 2009. Recruitment of functionally distinct membrane proteins to chromatin mediates nuclear envelope formation in vivo. The Journal of Cell Biology 186:183–191. doi:10.1083/jcb.200901106

Andrey P, Maurin Y. 2005. Free-D: an integrated environment for three-dimensional reconstruction from serial sections. J Neurosci Meth 145:233–244. doi:10.1016/j.jneumeth.2005.01.006

Andronov L, Ouararhni K, Stoll I, Klaholz BP, Hamiche A. 2019. CENP-A nucleosome clusters form rosette-like structures around HJURP during G1. Nat Commun 10:4436. doi:10.1038/s41467-019-12383-3

Appen A von, LaJoie D, Johnson IE, Trnka MJ, Pick SM, Burlingame AL, Ullman KS, Frost A. 2020. LEM2 phase separation governs ESCRT-mediated nuclear envelope reformation. Nature 582:115–118. doi:10.1038/s41586-020-2232-x

Arnoult N, Beneden AV, Decottignies A. 2012. Telomere length regulates TERRA levels through increased trimethylation of telomeric H3K9 and HP1α. doi:10.1038/nsmb.2364

Arnoult N, Schluth-Bolard C, Letessier A, Drascovic I, Bouarich-Bourimi R, Campisi J, Kim S-H, Boussouar A, Ottaviani A, Magdinier F, Gilson E, Londoño-Vallejo A. 2010. Replication timing of human telomeres is chromosome arm-specific, influenced by subtelomeric structures and connected to nuclear localization. PLoS genetics 6:e1000920. doi:10.1371/journal.pgen.1000920

Arpòn J, Sakai K, Gaudin V, Andrey P. 2021a. Spatial modeling of biological patterns shows multiscale organization of Arabidopsis thaliana heterochromatin. Nature Publishing Group 11:323. doi:10.1038/s41598-020-79158-5

Arpòn J, Sakai K, Gaudin V, Andrey P. 2021b. Spatial modeling of biological patterns shows multiscale organization of Arabidopsis thaliana heterochromatin. Sci Rep-uk 11:323. doi:10.1038/s41598-020-79158-5

Ball G, Demmerle J, Kaufmann R, Davis I, Dobbie IM, Schermelleh L. 2015a. SIMcheck: a Toolbox for Successful Super-resolution Structured Illumination Microscopy. Nature Publishing Group 5:15915. doi:10.1038/srep15915

Ball G, Demmerle J, Kaufmann R, Davis I, Dobbie IM, Schermelleh L. 2015b. SIMcheck: a Toolbox for Successful Super-resolution Structured Illumination Microscopy. Sci Rep-uk 5:15915. doi:10.1038/srep15915

Bickmore WA, Chubb JR. 2003. Chromosome Position: Now, Where Was I? Curr Biol 13:R357–R359. doi:10.1016/s0960-9822(03)00276-8

Bolzer A, Kreth G, Solovei I, Koehler D, Saracoglu K, Fauth C, Müller S, Eils R, Cremer C, Speicher MR, Cremer T. 2005. Three-Dimensional Maps of All Chromosomes in Human Male Fibroblast Nuclei and Prometaphase Rosettes. Plos Biol 3:e157. doi:10.1371/journal.pbio.0030157

Chikashige Y, Ding DQ, Funabiki H, Haraguchi T, Mashiko S, Yanagida M, Hiraoka Y. 1994. Telomere-led premeiotic chromosome movement in fission yeast. Science (New York, NY) 264:270–273. doi: 10.1126/science.8146661

Chikashige Y, Haraguchi T, Hiraoka Y. 2007. Another way to move chromosomes. Chromosoma 116:497–505. doi:10.1007/s00412-007-0114-8

Chikashige Y, Tsutsumi C, Yamane M, Okamasa K, Haraguchi T, Hiraoka Y. 2006. Meiotic proteins bqt1 and bqt2 tether telomeres to form the bouquet arrangement of chromosomes. Cell 125:59–69. doi:10.1016/j.cell.2006.01.048

Chikashige Y, Yamane M, Okamasa K, Tsutsumi C, Kojidani T, Sato M, Haraguchi T, Hiraoka Y. 2009. Membrane proteins Bqt3 and - 4 anchor telomeres to the nuclear envelope to ensure chromosomal bouquet formation. The Journal of Cell Biology 187:413–427. doi:10.1083/jcb.200902122

Chow TT, Shi X, Wei J-H, Guan J, Stadler G, Huang B, Blackburn EH. 2018. Local enrichment of HP1alpha at telomeres alters their structure and regulation of telomere protection. Nature communications 9:3583. doi:10.1038/s41467-018-05840-y

Chuang TCY, Moshir S, Garini Y, Chuang AY-C, Young IT, Vermolen B, Doel R van den, Mougey V, Perrin M, Braun M, Kerr PD, Fest T, Boukamp P, Mai S. 2004. The three-dimensional organization of telomeres in the nucleus of mammalian cells. BMC biology 2:12. doi:10.1186/1741-7007-2-12

Conrad MN, Lee C-Y, Wilkerson JL, Dresser ME. 2007. MPS3 mediates meiotic bouquet formation in Saccharomyces cerevisiae. Proceedings of the National Academy of Sciences of the United States of America 104:8863–8868. doi:10.1073/pnas.0606165104

Crabbe L, Cesare AJ, Kasuboski JM, Fitzpatrick JAJ, Karlseder J. 2012. Human Telomeres Are Tethered to the Nuclear Envelope during Postmitotic Nuclear Assembly. Cell reports. doi:10.1016/j.celrep.2012.11.019

Cremer M, Hase J von, Volm T, Brero A, Kreth G, Walter J, Fischer C, Solovei I, Cremer C, Cremer T. 2001. Non-random radial higher-order chromatin arrangements in nuclei of diploid human cells. Chromosome Res 9:541–567. doi:10.1023/a:1012495201697

Cremer T, Cremer M. 2010. Chromosome Territories. Csh Perspect Biol 2:a003889. doi:10.1101/cshperspect.a003889

Cubiles MD, Barroso S, Vaquero-Sedas MI, Enguix A, Aguilera A, Vega-Palas MA. 2018. Epigenetic features of human telomeres. Nucleic acids research 46:2347–2355. doi:10.1093/nar/gky006

Dechat T, Gajewski A, Korbei B, Gerlich D, Daigle N, Haraguchi T, Furukawa K, Ellenberg J, Foisner R. 2004. LAP2alpha and BAF transiently localize to telomeres and specific regions on chromatin during nuclear assembly. Journal of Cell Science 117:6117–6128. doi:10.1242/jcs.01529

Demmerle J, Innocent C, North AJ, Ball G, Müller M, Miron E, Matsuda A, Dobbie IM, Markaki Y, Schermelleh L. 2017. Strategic and practical guidelines for successful structured illumination microscopy. Nat Protoc 12:988–1010. doi:10.1038/nprot.2017.019

Essers J, Cappellen WA van, Theil AF, Drunen E van, Jaspers NGJ, Hoeijmakers JHJ, Wyman C, Vermeulen W, Kanaar R. 2005. Dynamics of Relative Chromosome Position during the Cell Cycle. Mol Biol Cell 16:769–775. doi:10.1091/mbc.e04-10-0876

Falk M, Feodorova Y, Naumova N, Imakaev M, Lajoie BR, Leonhardt H, Joffe B, Dekker J, Fudenberg G, Solovei I, Mirny LA. 2019. Heterochromatin drives compartmentalization of inverted and conventional nuclei. Nature 570:395–399. doi:10.1038/s41586-019-1275-3

Funabiki H, Hagan I, Uzawa S, Yanagida M. 1993. Cell cycle-dependent specific positioning and clustering of centromeres and telomeres in fission yeast. The Journal of Cell Biology 121:961–976. doi:10.1083/jcb.121.5.961

Gerlich D, Beaudouin J, Kalbfuss B, Daigle N, Eils R, Ellenberg J. 2003. Global chromosome positions are transmitted through mitosis in mammalian cells. Cell 112:751–764.

Gruenbaum Y, Foisner R. 2015. Lamins: nuclear intermediate filament proteins with fundamental functions in nuclear mechanics and genome regulation. Annu Rev Biochem 84:131–164. doi:10.1146/annurev-biochem-060614-034115

Guacci V, Hogan E, Koshland D. 1997. Centromere position in budding yeast: evidence for anaphase A. Molecular biology of the cell 8:957–972. doi:10.1091/mbc.8.6.957

Guelen L, Pagie L, Brasset E, Meuleman W, Faza MB, Talhout W, Eussen BH, Klein A de, Wessels L, Laat WD, Steensel B van. 2008. Domain organization of human chromosomes revealed by mapping of nuclear lamina interactions. Nature 453:948–951. doi:10.1038/nature06947

Güttinger S, Laurell E, Kutay U. 2009. Orchestrating nuclear envelope disassembly and reassembly during mitosis. Nature reviews Molecular cell biology 10:178–191. doi:10.1038/nrm2641

Haraguchi T, Kojidani T, Koujin T, Shimi T, Osakada H, Mori C, Yamamoto A, Hiraoka Y. 2008. Live cell imaging and electron microscopy reveal dynamic processes of BAF-directed nuclear envelope assembly. Journal of Cell Science 121:2540–2554. doi:10.1242/jcs.033597

Haraguchi T, Koujin T, Hayakawa T, Kaneda T, Tsutsumi C, Imamoto N, Akazawa C, Sukegawa J, Yoneda Y, Hiraoka Y. 2000. Live fluorescence imaging reveals early recruitment of emerin, LBR, RanBP2, and Nup153 to reforming functional nuclear envelopes. Journal of Cell Science 113 ( Pt 5):779–794.

Hediger F, Neumann FR, Houwe GV, Dubrana K, Gasser SM. 2002. Live imaging of telomeres: yKu and Sir proteins define redundant telomere-anchoring pathways in yeast. Current biology: CB 12:2076–2089.

Jin QW, Fuchs J, Loidl J. 2000. Centromere clustering is a major determinant of yeast interphase nuclear organization. Journal of Cell Science 113 ( Pt 11):1903–1912.

Kaya-Okur HS, Wu SJ, Codomo CA, Pledger ES, Bryson TD, Henikoff JG, Ahmad K, Henikoff S. 2019. CUT&Tag for efficient epigenomic profiling of small samples and single cells. Nat Commun 10:1930. doi:10.1038/s41467-019-09982-5

Kind J, Pagie L, Ortabozkoyun H, Boyle S, Vries SS de, Janssen H, Amendola M, Nolen LD, Bickmore WA, Steensel B van. 2013. Single-cell dynamics of genome-nuclear lamina interactions. Cell 153:178–192. doi:10.1016/j.cell.2013.02.028

Kind J, Pagie L, Vries SS de, Nahidiazar L, Dey SS, Bienko M, Zhan Y, Lajoie B, Graaf CA de, Amendola M, Fudenberg G, Imakaev M, Mirny LA, Jalink K, Dekker J, Oudenaarden A van, Steensel B van. 2015. Genome-wide Maps of Nuclear Lamina Interactions in Single Human Cells. Cell 1–15. doi:10.1016/j.cell.2015.08.040

Kind J, Steensel B van. 2014. Stochastic genome-nuclear lamina interactions: modulating roles of Lamin A and BAF. Nucleus (Austin, Tex) 5:124–130. doi:10.4161/nucl.28825

Kraus F, Miron E, Demmerle J, Chitiashvili T, Budco A, Alle Q, Matsuda A, Leonhardt H, Schermelleh L, Markaki Y. 2017. Quantitative 3D structured illumination microscopy of nuclear structures. Nat Protoc 12:1011–1028. doi:10.1038/nprot.2017.020

Kutay U, Hetzer MW. 2008. Reorganization of the nuclear envelope during open mitosis. Current opinion in cell biology 20:669–677. doi:10.1016/j.ceb.2008.09.010

Kychygina A, Dall’Osto M, Allen JAM, Cadoret J-C, Piras V, Pickett HA, Crabbe L. 2021a. Progerin impairs 3D genome organization and induces fragile telomeres by limiting the dNTP pools. Sci Rep-uk 11:13195. doi:10.1038/s41598-021-92631-z

Kychygina A, Dall’Osto M, Allen JAM, Cadoret J-C, Piras V, Pickett HA, Crabbe L. 2021b. Progerin impairs 3D genome organization and induces fragile telomeres by limiting the dNTP pools. Sci Rep-uk 11:13195. doi:10.1038/s41598-021-92631-z

Lawo S, Hasegan M, Gupta GD, Pelletier L. 2012. Subdiffraction imaging of centrosomes reveals higher-order organizational features of pericentriolar material. Nat Cell Biol 14:1148–1158. doi:10.1038/ncb2591

Lemaitre C, Fischer B, Kalousi A, Hoffbeck A-S, Guirouilh-Barbat J, Shahar OD, Genet D, Goldberg M, Betrand P, Lopez B, Brino L, Soutoglou E. 2012. The nucleoporin 153, a novel factor in double-strand break repair and DNA damage response. 31:4803–4809. doi:10.1038/onc.2011.638

Link J, Jahn D, Alsheimer M. 2015. Structural and functional adaptations of the mammalian nuclear envelope to meet the meiotic requirements. Nucleus (Austin, Tex) 6:93–101. doi:10.1080/19491034.2015.1004941

Lorensen WE, Cline HE. 1987. Marching cubes: A high resolution 3D surface construction algorithm. Acm Siggraph Comput Graph 21:163–169. doi:10.1145/37401.37422

Martin C, Chen S, Jackson DA. 2010. Inheriting nuclear organization: can nuclear lamins impart spatial memory during post-mitotic nuclear assembly? Chromosome research: an international journal on the molecular, supramolecular and evolutionary aspects of chromosome biology 18:525–541. doi:10.1007/s10577-010-9137-8

Mayer R, Brero A, Hase J von, Schroeder T, Cremer T, Dietzel S. 2005. Common themes and cell type specific variations of higher order chromatin arrangements in the mouse. Bmc Cell Biol 6:44–44. doi:10.1186/1471-2121-6-44

Molenaar C, Wiesmeijer K, Verwoerd NP, Khazen S, Eils R, Tanke HJ, Dirks RW. 2003. Visualizing telomere dynamics in living mammalian cells using PNA probes. The EMBO Journal 22:6631–6641. doi:10.1093/emboj/cdg633

Muller H, Jr JG, Drinnenberg IA. 2019. The Impact of Centromeres on Spatial Genome Architecture. Trends Genet 35:565–578. doi:10.1016/j.tig.2019.05.003

Müller I, Boyle S, Singer RH, Bickmore WA, Chubb JR. 2010. Stable Morphology, but Dynamic Internal Reorganisation, of Interphase Human Chromosomes in Living Cells. Plos One 5:e11560. doi:10.1371/journal.pone.0011560

Nagele RG, Velasco AQ, Anderson WJ, McMahon DJ, Thomson Z, Fazekas J, Wind K, Lee H. 2001. Telomere associations in interphase nuclei: possible role in maintenance of interphase chromosome topology. J Cell Sci 114:377–388. doi:10.1242/jcs.114.2.377

Naumova N, Imakaev M, Fudenberg G, Zhan Y, Lajoie BR, Mirny LA, Dekker J. 2013. Organization of the mitotic chromosome. Science (New York, NY) 342:948–953. doi:10.1126/science.1236083

Olins AL, Rhodes G, Welch DBM, Zwerger M, Olins DE. 2010. Lamin B receptor: Multi-tasking at the nuclear envelope. Nucleus (Austin, Tex) 1:53–70. doi:10.4161/nucl.1.1.10515

Olmos Y, Hodgson L, Mantell J, Verkade P, Carlton JG. 2015. ESCRT-III controls nuclear envelope reformation. Nature 522:236–239. doi:10.1038/nature14503

O’Sullivan RJ, Kubicek S, Schreiber SL, Karlseder J. 2010. Reduced histone biosynthesis and chromatin changes arising from a damage signal at telomeres. 17:1218–1225. doi:10.1038/nsmb.1897

Park S-Y, Park J-E, Kim T-S, Kim JH, Kwak M-J, Ku B, Tian L, Murugan RN, Ahn M, Komiya S, Hojo H, Kim N-H, Kim BY, Bang JK, Erikson RL, Lee KW, Kim SJ, Oh B-H, Yang W, Lee KS. 2014. Molecular Basis for Unidirectional Scaffold Switching of Human Plk4 in Centriole Biogenesis. Nat Struct Mol Biol 21:696–703. doi:10.1038/nsmb.2846

Peric-Hupkes D, Meuleman W, Pagie L, Bruggeman SWM, Solovei I, Brugman W, Gräf S, Flicek P, Kerkhoven RM, Lohuizen M van, Reinders M, Wessels L, Steensel B van. 2010. Molecular maps of the reorganization of genome-nuclear lamina interactions during differentiation. Molecular Cell 38:603–613. doi:10.1016/j.molcel.2010.03.016

Piel M, Meyer P, Khodjakov A, Rieder CL, Bornens M. 2000. The Respective Contributions of the Mother and Daughter Centrioles to Centrosome Activity and Behavior in Vertebrate Cells. J Cell Biology 149:317–330. doi:10.1083/jcb.149.2.317

Poleshko A, Mansfield KM, Burlingame CC, Andrake MD, Shah NR, Katz RA. 2013. The Human Protein PRR14 Tethers Heterochromatin to the Nuclear Lamina during Interphase and Mitotic Exit. Cell reports 5:292–301. doi:10.1016/j.celrep.2013.09.024

Poleshko A, Smith CL, Nguyen SC, Sivaramakrishnan P, Wong KG, Murray JI, Lakadamyali M, Joyce EF, Jain R, Epstein JA. 2019a. H3K9me2 orchestrates inheritance of spatial positioning of peripheral heterochromatin through mitosis. Elife 8:e49278. doi:10.7554/elife.49278

Poleshko A, Smith CL, Nguyen SC, Sivaramakrishnan P, Wong KG, Murray JI, Lakadamyali M, Joyce EF, Jain R, Epstein JA. 2019b. H3K9me2 orchestrates inheritance of spatial positioning of peripheral heterochromatin through mitosis. eLife 8:1–24. doi:10.7554/elife.49278

Postma M, Goedhart J. 2019. PlotsOfData—A web app for visualizing data together with their summaries. Plos Biol 17:e3000202. doi:10.1371/journal.pbio.3000202

Ptak C, Wozniak RW. 2016. ScienceDirect Nucleoporins and chromatin metabolism. Current opinion in cell biology 40:153–160. doi:10.1016/j.ceb.2016.03.024

Samwer M, Schneider MWG, Hoefler R, Schmalhorst PS, Jude JG, Zuber J, Gerlich DW. 2017. DNA Cross-Bridging Shapes a Single Nucleus from a Set of Mitotic Chromosomes 1-41. doi:10.1016/j.cell.2017.07.038

Schaik T van, Vos M, Peric-Hupkes D, Celie PH, Steensel B van. 2020. Cell cycle dynamics of lamina-associated DNA. Embo Rep 21:e50636. doi:10.15252/embr.202050636

Schermelleh L, Ferrand A, Huser T, Eggeling C, Sauer M, Biehlmaier O, Drummen GPC. 2019. Superresolution microscopy demystified. Nature Cell Biology 21:72–84. doi:10.1038/s41556-018-0251-8

Schindelin J, Arganda-Carreras I, Frise E, Kaynig V, Longair M, Pietzsch T, Preibisch S, Rueden C, Saalfeld S, Schmid B, Tinevez J-Y, White DJ, Hartenstein V, Eliceiri K, Tomancak P, Cardona A. 2012. Fiji: an open-source platform for biological-image analysis. Nat Methods 9:676–682. doi:10.1038/nmeth.2019

Schmälter A-K, Kuzyk A, Righolt CH, Neusser M, Steinlein OK, Müller S, Mai S. 2014. Distinct nuclear orientation patterns for mouse chromosome 11 in normal B lymphocytes. Bmc Cell Biol 15:22–22. doi:10.1186/1471-2121-15-22

Schober H, Ferreira H, Kalck V, Gehlen LR, Gasser SM. 2009. Yeast telomerase and the SUN domain protein Mps3 anchor telomeres and repress subtelomeric recombination. Genes & Development 23:928–938. doi:10.1101/gad.1787509

Sehgal N, Fritz AJ, Vecerova J, Ding H, Chen Z, Stojkovic B, Bhattacharya S, Xu J, Berezney R. 2016. Large-scale probabilistic 3D organization of human chromosome territories. Hum Mol Genet 25:419–436. doi:10.1093/hmg/ddv479

Shevelyov YY. 2020. The Role of Nucleoporin Elys in Nuclear Pore Complex Assembly and Regulation of Genome Architecture. Int J Mol Sci 21:9475. doi:10.3390/ijms21249475

Sobecki M, Souaid C, Boulay J, Guerineau V, Noordermeer D, Crabbe L. 2018. MadID, a Versatile Approach to Map Protein-DNA Interactions, Highlights Telomere-Nuclear Envelope Contact Sites in Human Cells. Cell Reports 25:2891–2903.e5. doi:10.1016/j.celrep.2018.11.027

Soille P. 2004. Morphological Image Analysis, Principles and Applications 241–265. doi:10.1007/978-3-662-05088-0_8

Solovei I, Schermelleh L, Düring K, Engelhardt A, Stein S, Cremer C, Cremer T. 2004. Differences in centromere positioning of cycling and postmitotic human cell types. Chromosoma 112:410–423. doi:10.1007/s00412-004-0287-3

Solovei I, Thanisch K, Feodorova Y. 2016. How to rule the nucleus: divide et impera. Current opinion in cell biology 40:47–59. doi:10.1016/j.ceb.2016.02.014

Solovei I, Wang AS, Thanisch K, Schmidt CS, Krebs S, Zwerger M, Cohen TV, Devys D, Foisner R, Peichl L, Herrmann H, Blum H, Engelkamp D, Stewart CL, Leonhardt H, Joffe B. 2013. LBR and lamin A/C sequentially tether peripheral heterochromatin and inversely regulate differentiation. Cell 152:584–598. doi:10.1016/j.cell.2013.01.009

Stirling DR, Swain-Bowden MJ, Lucas AM, Carpenter AE, Cimini BA, Goodman A. 2021. CellProfiler 4: improvements in speed, utility and usability. Bmc Bioinformatics 22:433. doi:10.1186/s12859-021-04344-9

Szczurek A, Contu F, Hoang A, Dobrucki J, Mai S. 2018a. Aqueous mounting media increasing tissue translucence improve image quality in Structured Illumination Microscopy of thick biological specimen. Nature Publishing Group 8:13971. doi:10.1038/s41598-018-32191-x

Szczurek A, Contu F, Hoang A, Dobrucki J, Mai S. 2018b. Aqueous mounting media increasing tissue translucence improve image quality in Structured Illumination Microscopy of thick biological specimen. Sci Rep-uk 8:13971. doi:10.1038/s41598-018-32191-x

Talamas JA, Hetzer MW. 2011. POM121 and Sun1 play a role in early steps of interphase NPC assembly. The Journal of Cell Biology 194:27–37. doi:10.1083/jcb.201012154

Thomson I, Gilchrist S, Bickmore WA, Chubb JR. 2004. The radial positioning of chromatin is not inherited through mitosis but is established de novo in early G1. Current biology: CB 14:166–172.

Umlauf D, Sobecki M, Crabbe L. 2020. Methyl Adenine Identification (MadID): High-Resolution Detection of Protein-DNA Interactions. Methods Mol Biology Clifton N J 2175:123–138. doi: 10.1007/978-1-0716-0763-3_10

Vietri M, Schink KO, Campsteijn C, Wegner CS, Schultz SW, Christ L, Thoresen SB, Brech A, Raiborg C, Stenmark H. 2015. Spastin and ESCRT-III coordinate mitotic spindle disassembly and nuclear envelope sealing. Nature 522:231–235. doi:10.1038/nature14408

Vincent L, Soille P. 1991. Watersheds in digital spaces: an efficient algorithm based on immersion simulations. Ieee T Pattern Anal 13:583–598. doi:10.1109/34.87344

Vourc’h C, Taruscio D, Boyle AL, Ward DC. 1993. Cell cycle-dependent distribution of telomeres, centromeres, and chromosome-specific subsatellite domains in the interphase nucleus of mouse lymphocytes. Experimental cell research 205:142–151. doi:10.1006/excr.1993.1068

Walter J, Schermelleh L, Cremer M, Tashiro S, Cremer T. 2003a. Chromosome order in HeLa cells changes during mitosis and early G1, but is stably maintained during subsequent interphase stages. J Cell Biology 160:685–697. doi:10.1083/jcb.200211103

Walter J, Schermelleh L, Cremer M, Tashiro S, Cremer T. 2003b. Chromosome order in HeLa cells changes during mitosis and early G1, but is stably maintained during subsequent interphase stages. The Journal of Cell Biology 160:685–697. doi:10.1083/jcb.200211103

Weierich C, Brero A, Stein S, Hase J von, Cremer C, Cremer T, Solovei I. 2003. Three-dimensional arrangements of centromeres and telomeres in nuclei of human and murine lymphocytes. Chromosome research: an international journal on the molecular, supramolecular and evolutionary aspects of chromosome biology 11:485–502.

Williams RRE, Fisher AG. 2003. Chromosomes, positions please! Nat Cell Biol 5:388–390. doi:10.1038/ncb0503-388

Zullo JM, Demarco IA, Piqué-Regi R, Gaffney DJ, Epstein CB, Spooner CJ, Luperchio TR, Bernstein BE, Pritchard JK, Reddy KL, Singh H. 2012. DNA sequence-dependent compartmentalization and silencing of chromatin at the nuclear lamina. Cell 149:1474–1487. doi:10.1016/j.cell.2012.04.035

