## Supplementary figures and images for "Spatial modeling of telomere intra-nuclear distribution reveals non-random organization that varies during cell cycle and depends on LAP2 and BAF"

### Supplemental figures

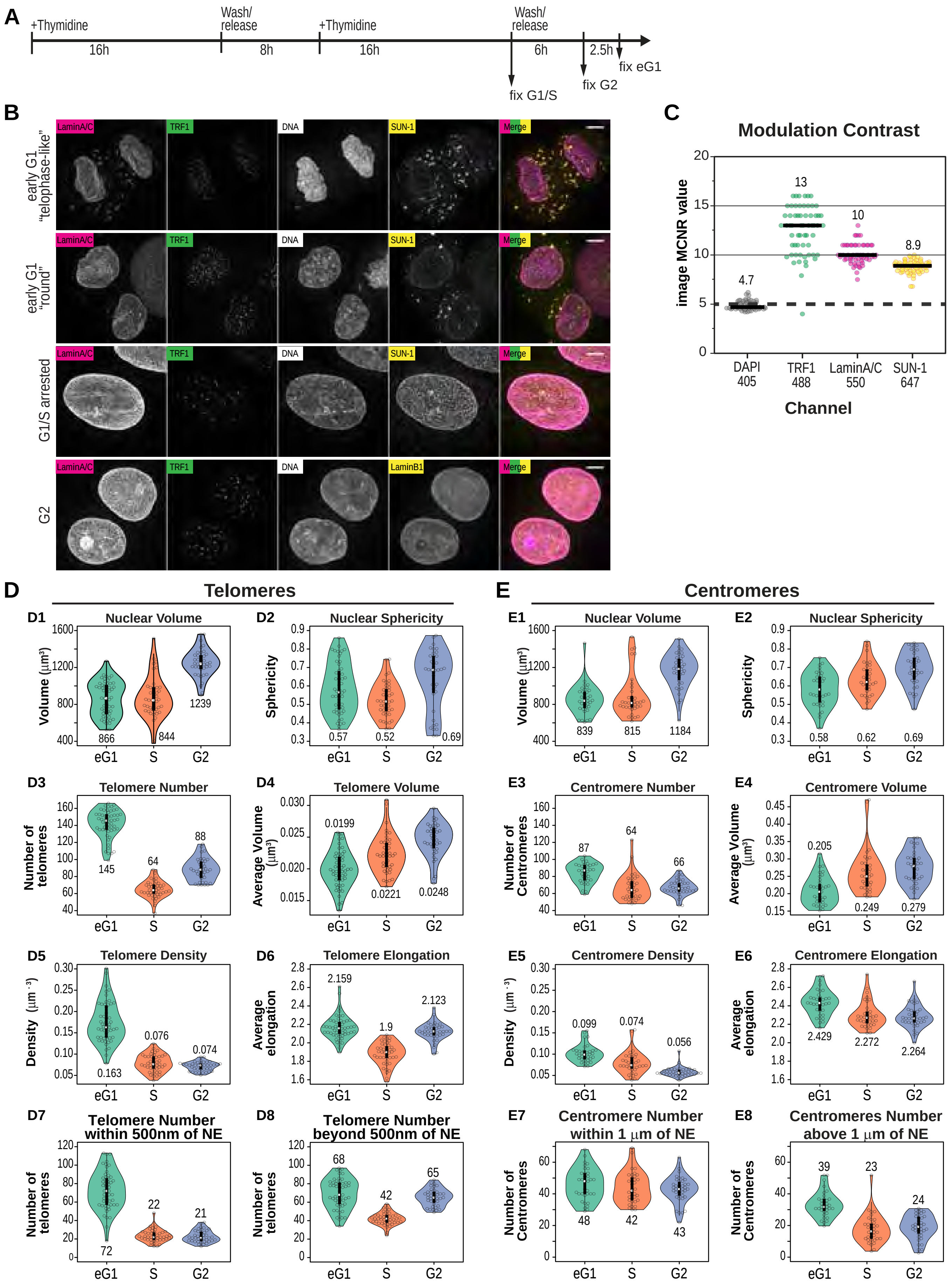

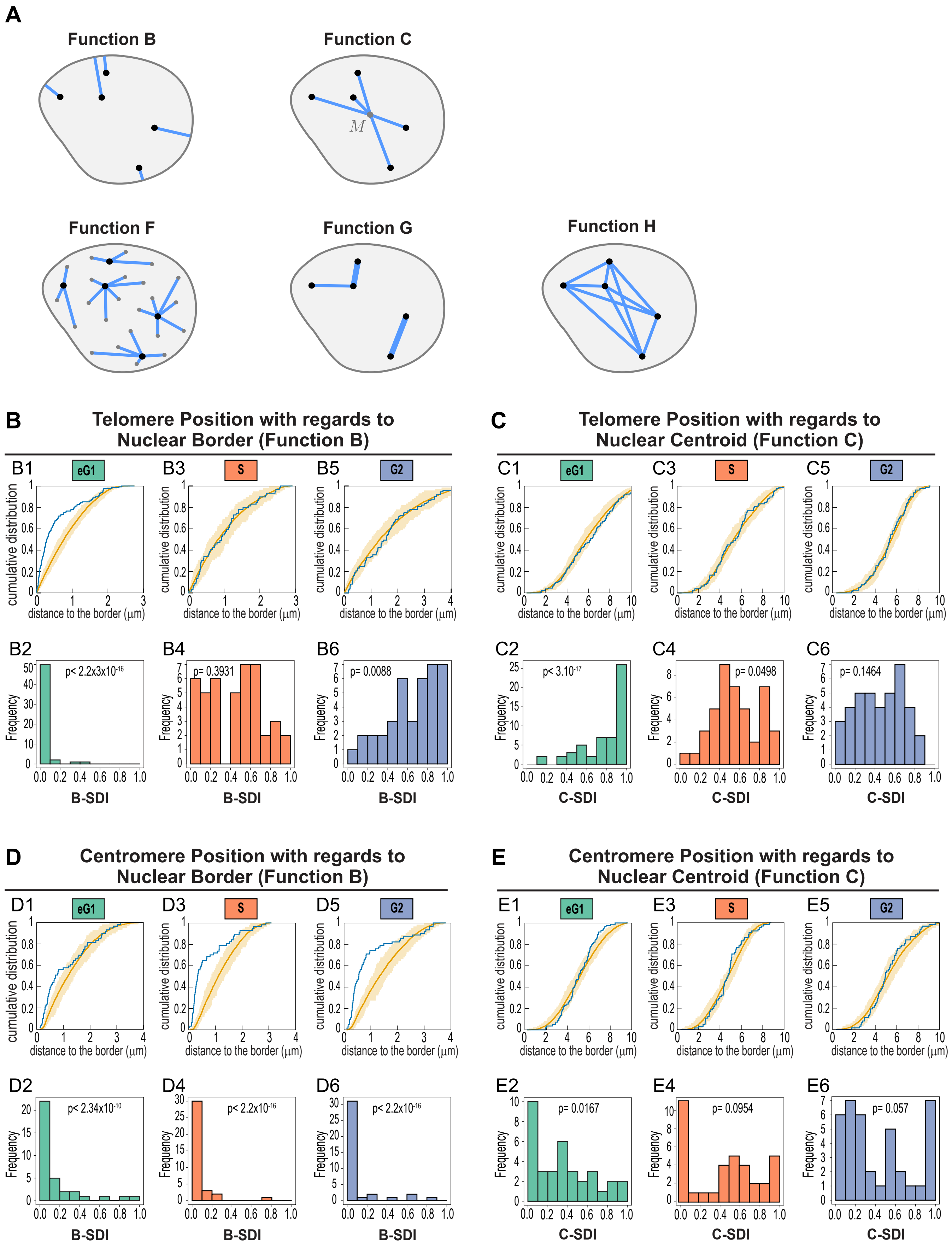

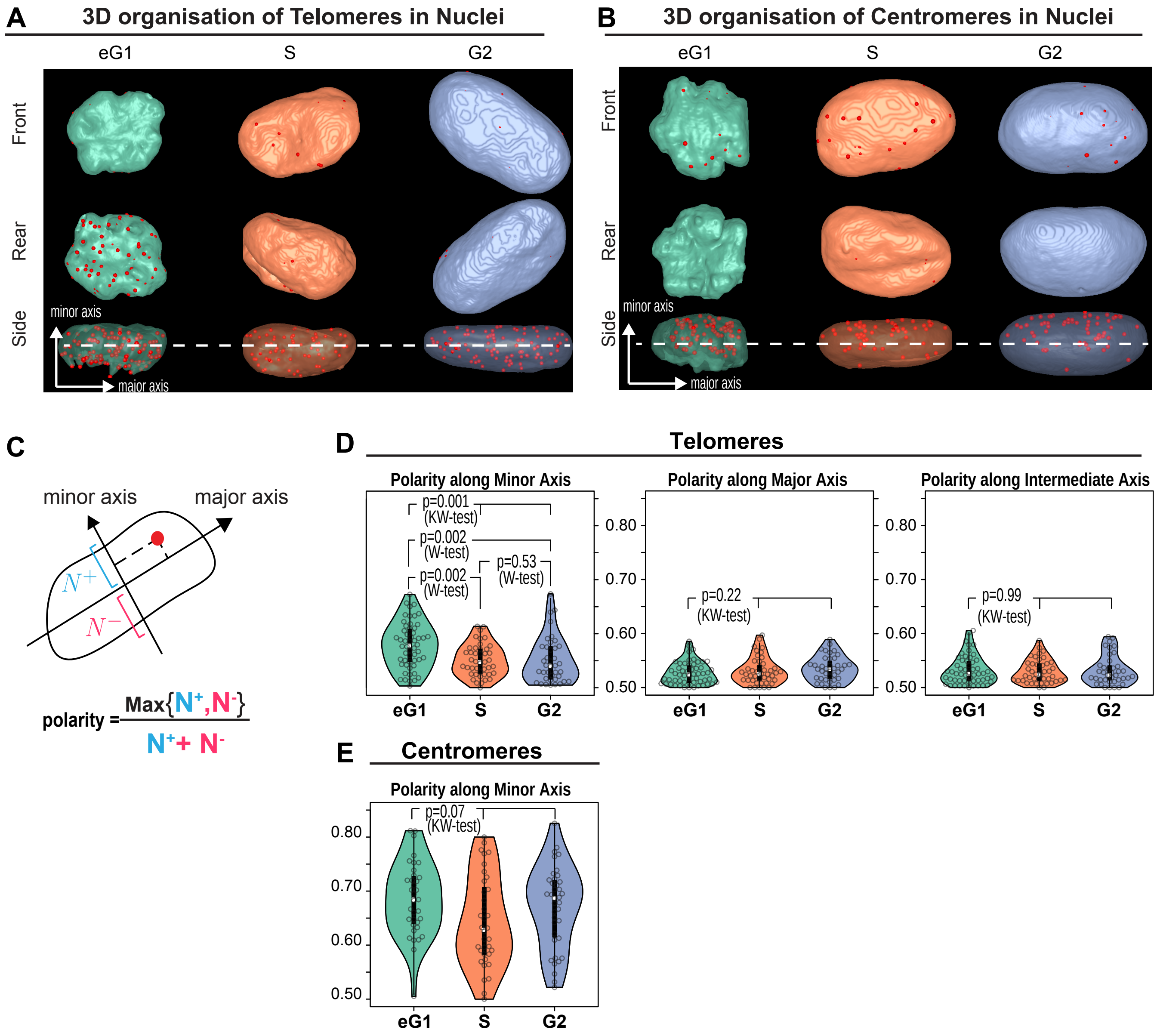

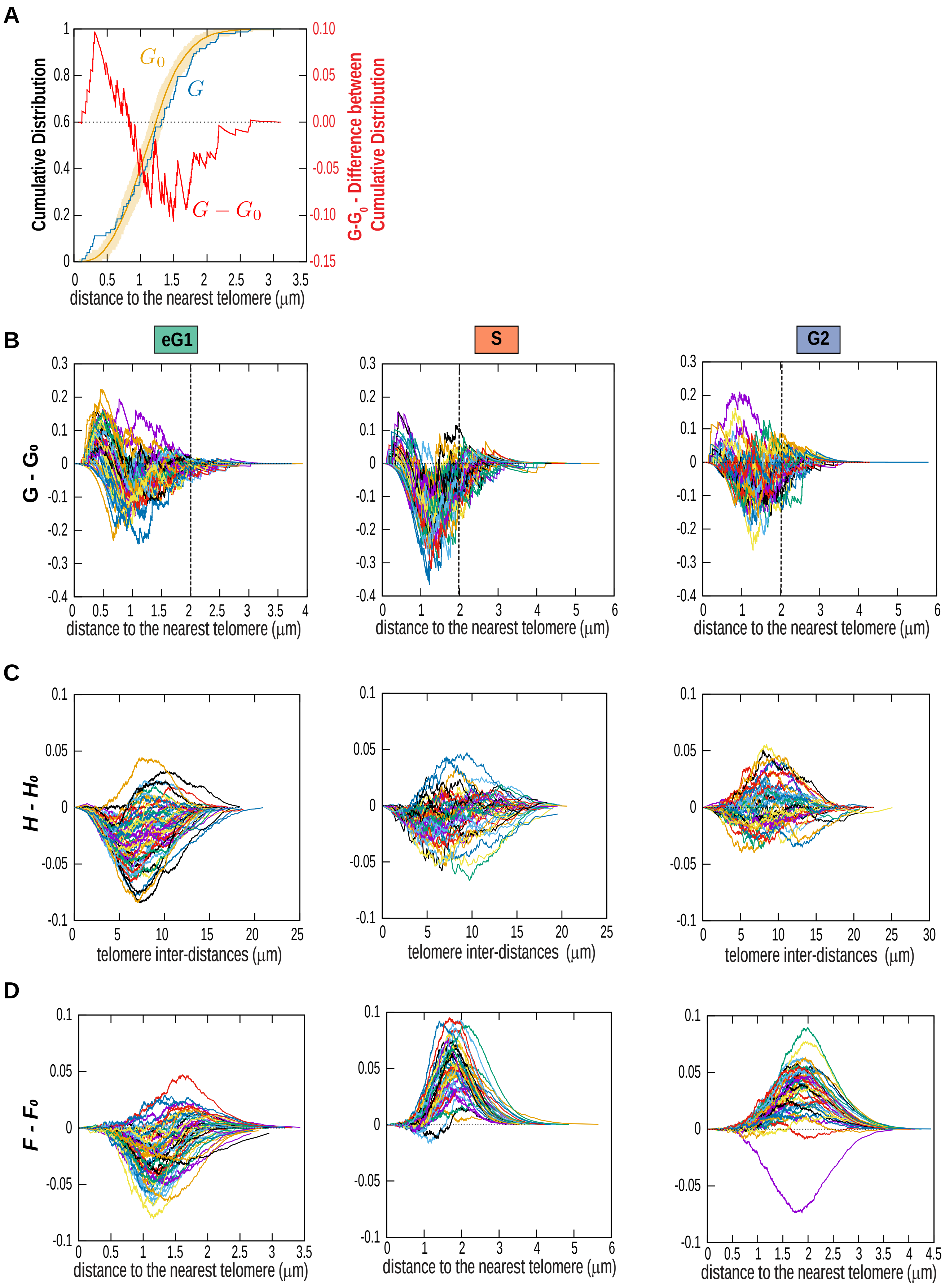

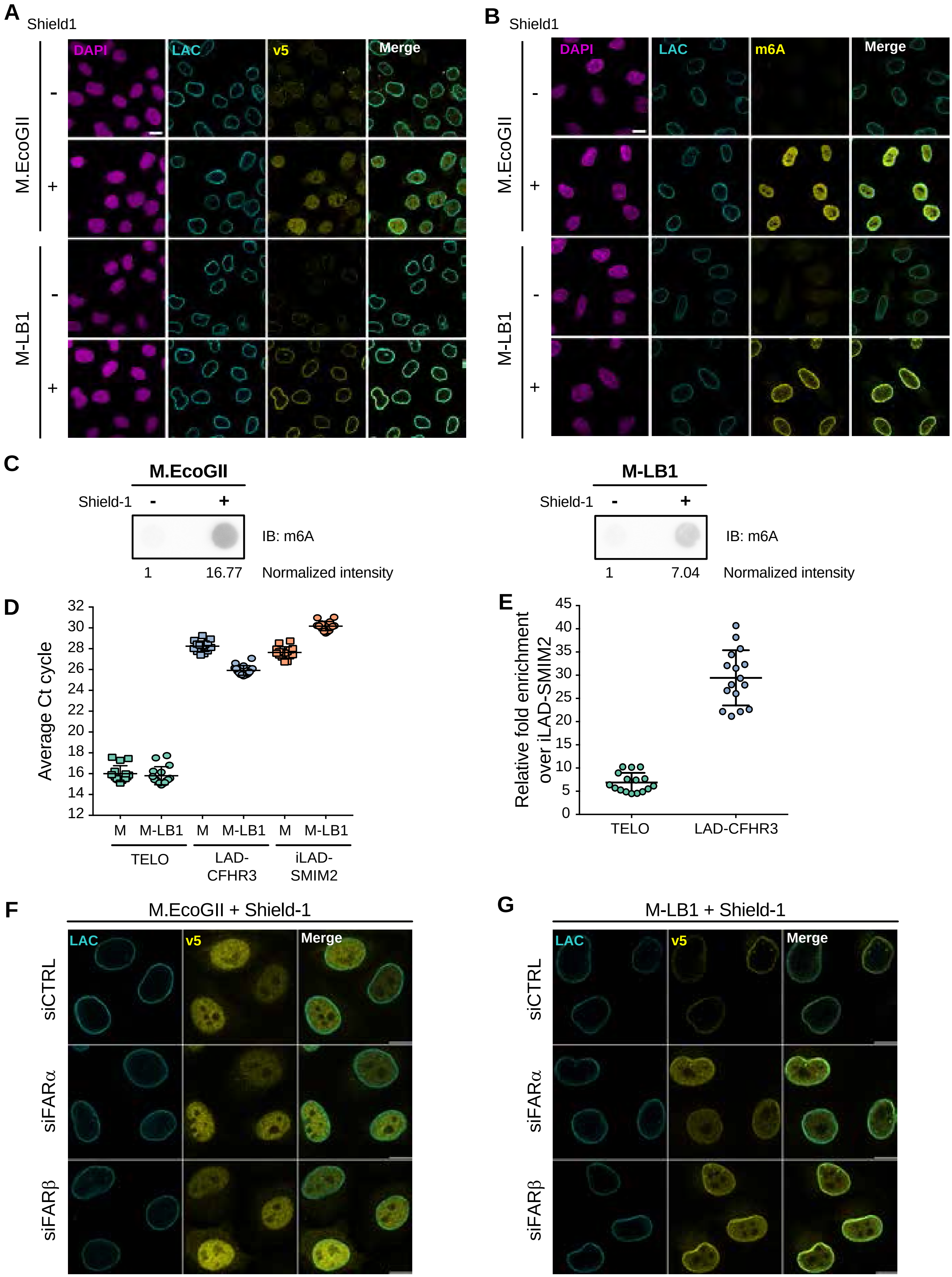

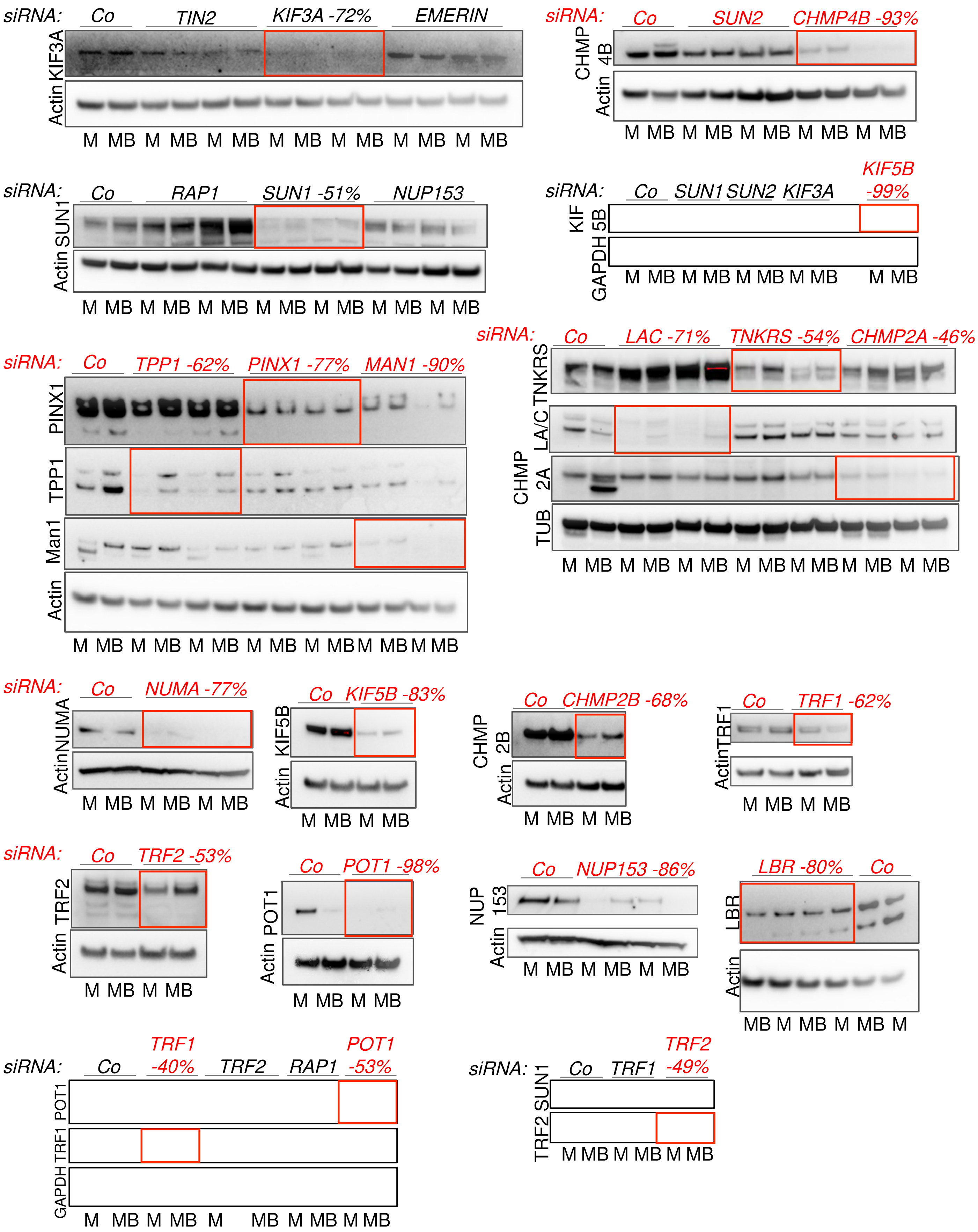

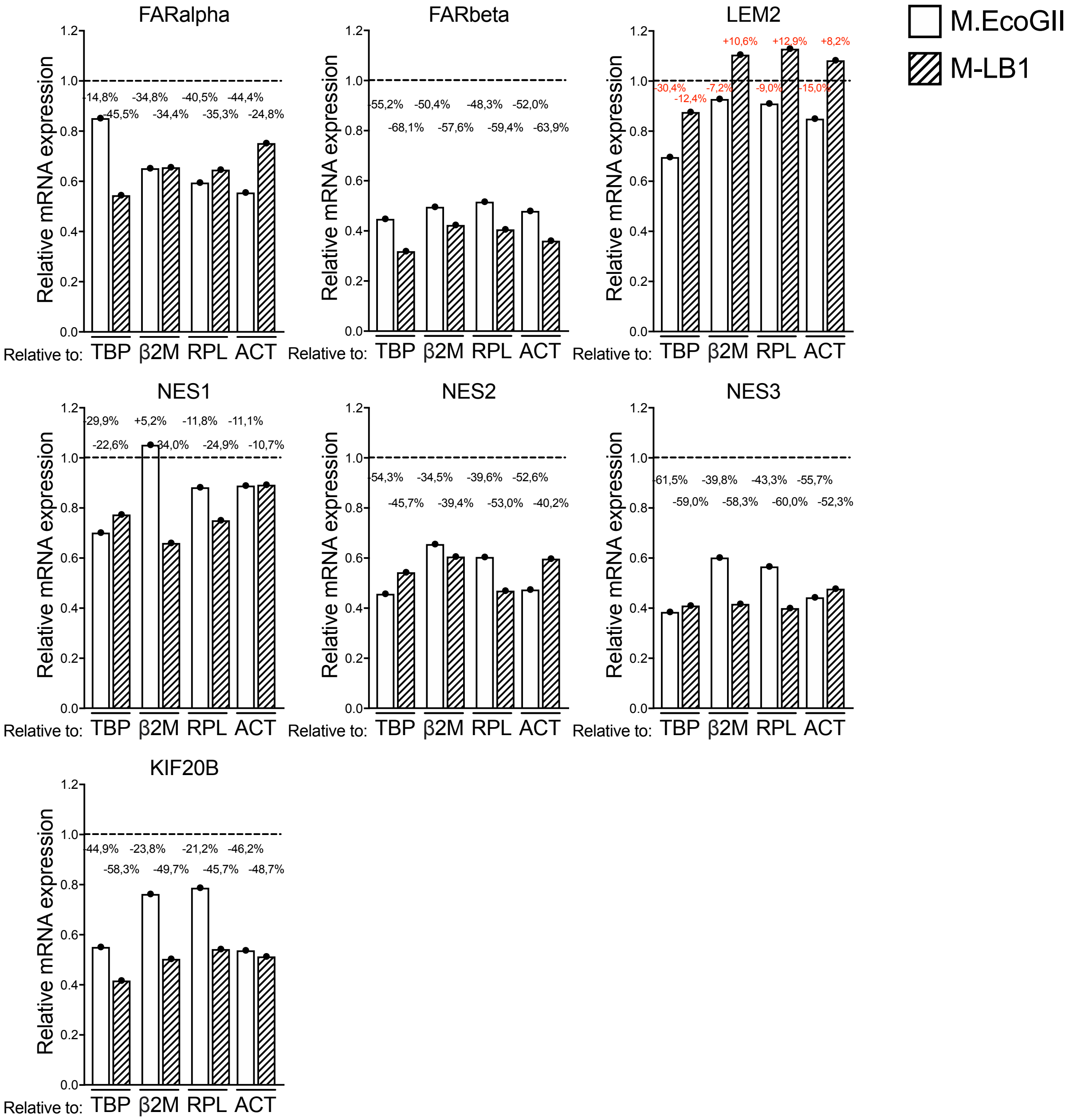

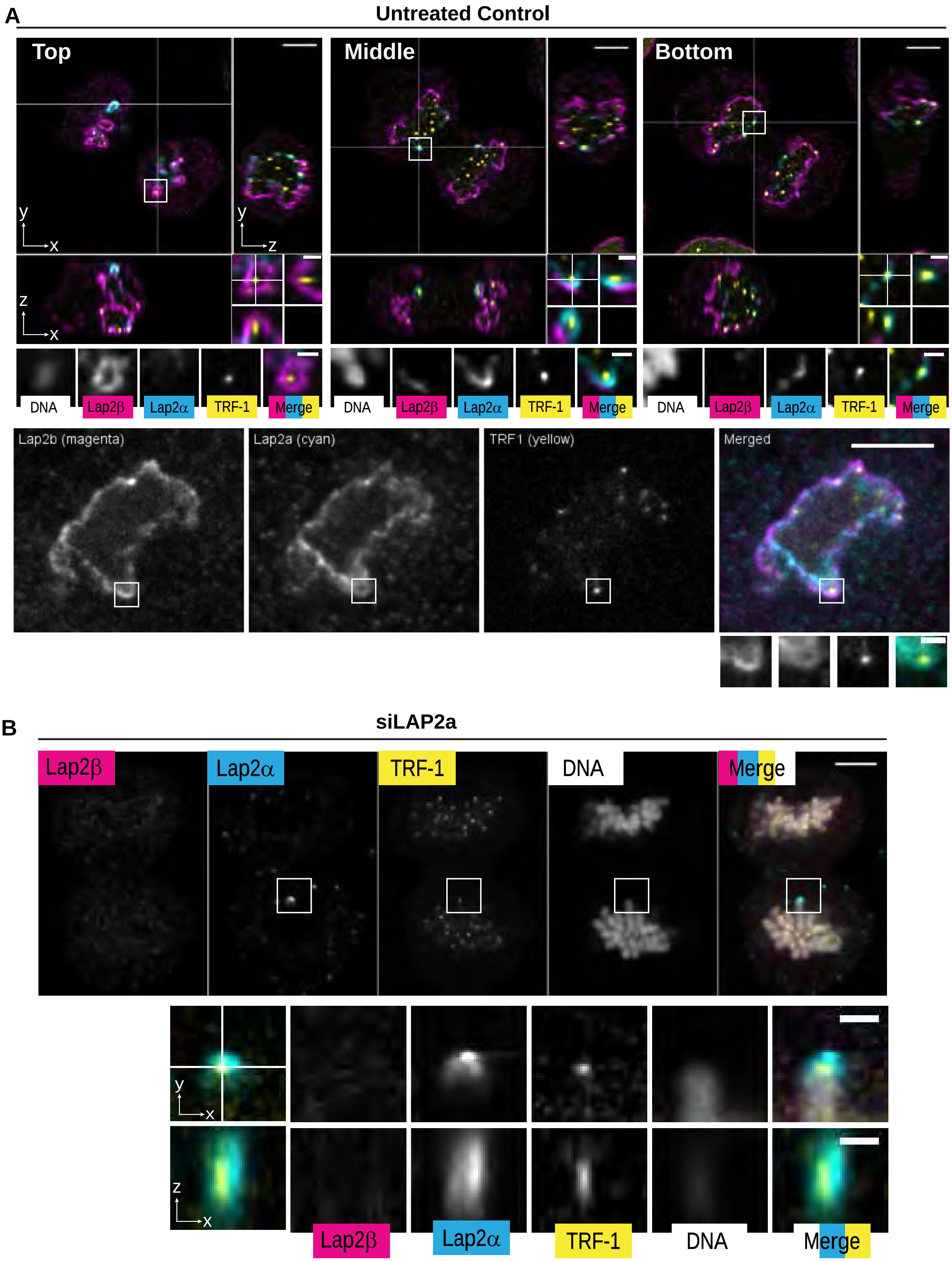

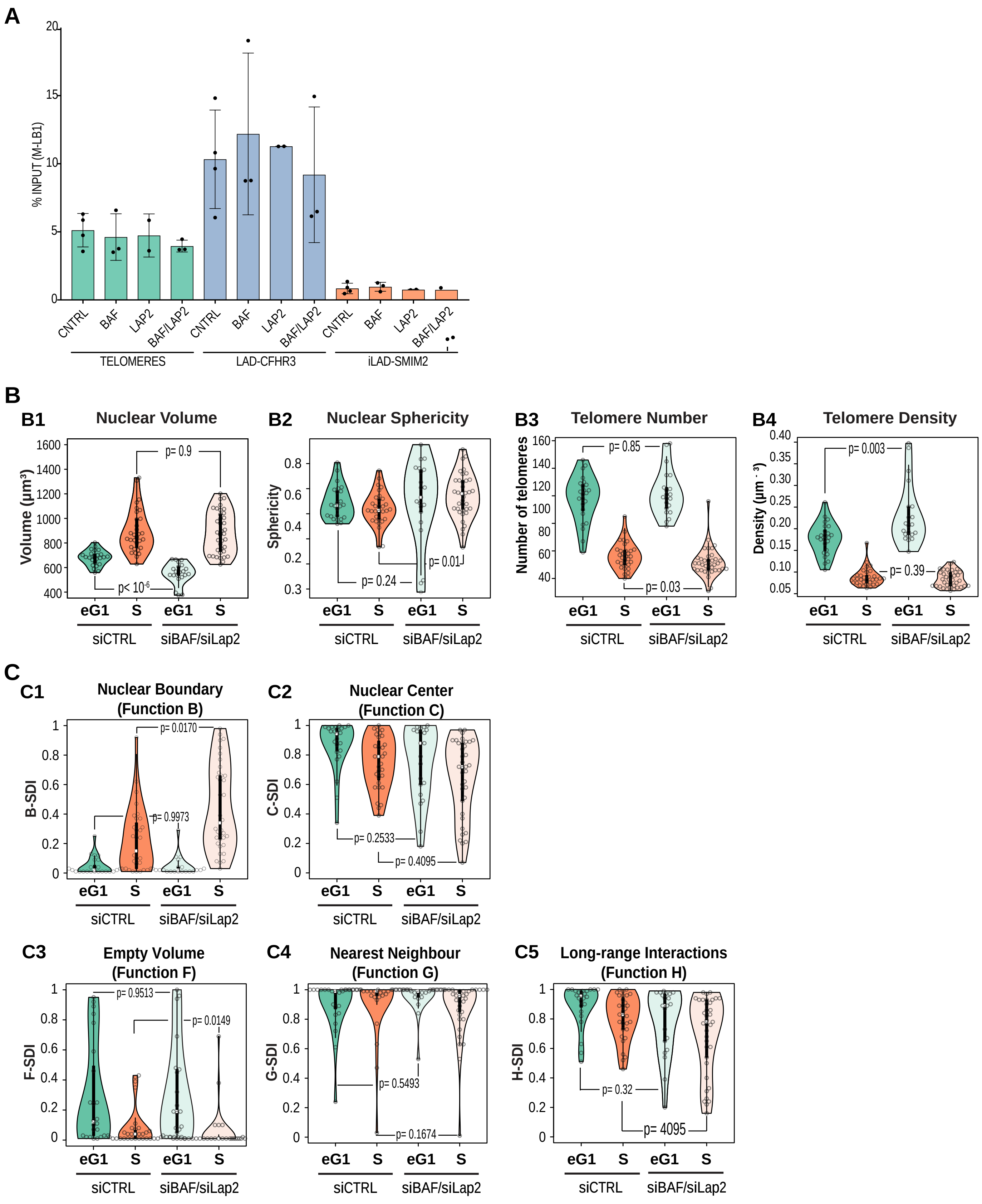
